# The EZH2 inhibitor tazemetostat mitigates HIV immune evasion, reduces reservoir formation, and promotes durable CD8⁺ T-cell revitalization

**DOI:** 10.1101/2024.10.11.617869

**Authors:** Andrea Gramatica, Itzayana G. Miller, Adam R. Ward, Farzana Khan, Tyler J. Kemmer, Jared Weiler, Tan Thinh Huynh, Paul Zumbo, Andrew P. Kurland, Louise Leyre, Yanqin Ren, Thais Klevorn, Dennis C. Copertino, Uchenna Chukwukere, Callie Levinger, Thomas R. Dilling, Noemi Linden, Nathan L. Board, Emma Falling Iversen, Sandra Terry, Talia M. Mota, Seden Bedir, Kiera L. Clayton, Alberto Bosque, Lynsay MacLaren Ehui, Colin Kovacs, Doron Betel, Jeffry R. Johnson, Mirko Paiardini, Ali Danesh, R. Brad Jones

## Abstract

Persistent HIV reservoirs in CD4⁺ T-cells pose a barrier to curing HIV infection. We identified overexpression of enhancer of zeste homolog 2 (EZH2) in HIV-infected CD4⁺ T-cells that survive cytotoxic T lymphocyte (CTL) exposure, suggesting a mechanism of CTL resistance. Inhibition of EZH2 with the FDA-approved drug tazemetostat increased surface expression of major histocompatibility complex class I (MHC-I) on CD4⁺ T-cells, counterbalancing HIV Nef–mediated MHC-I downregulation. This improved CTL-mediated elimination of HIV-infected cells and suppressed viral replication in vitro. In a participant-derived xenograft mouse model, tazemetostat elevated MHC-I and the pro-apoptotic protein BIM in CD4⁺ T-cells, facilitating CD8⁺ T-cell–mediated reductions of HIV reservoir seeding. Additionally, tazemetostat promoted sustained skewing of CD8⁺ T-cells toward less differentiated and exhausted phenotypes. Our findings reveal EZH2 overexpression as a novel mechanism of CTL resistance and support the clinical evaluation of tazemetostat to enhance clearance of HIV reservoirs and improve CD8+ T-cell function.

## Introduction

HIV replication can be suppressed by antiretroviral therapy (ART), preventing disease progression and viral transmission. However, a safe and scalable cure remains elusive due to the persistence of viral reservoirs, primarily CD4^+^ T-cells harboring integrated HIV proviruses. These reservoirs evade immune elimination largely through viral latency and reseed viral replication if ART is discontinued^1–3^. Cure strategies aim for either ‘ART-free virologic control’ or complete viral eradication^4,5^.

HIV-specific CD8^+^ T-cell responses hold promise for contributing to both goals^6–8^. In the context of ART-free virologic control, these cells play a critical role in “elite controllers,” individuals who control HIV replication to undetectable levels without ART^9–14^. For viral eradication, leveraging the specific recognition and elimination of infected cells by CD8^+^ cytotoxic T-lymphocyte (CTL) T-cells will likely be necessary. While harnessing CD8^+^ T-cells for either strategy has been challenging, recent clinical trial results and novel insights provide both optimism and direction for ongoing efforts^15^.

The period of ART initiation represents a window of opportunity for immune-based interventions that can durably impact virologic outcomes, given that much of the HIV reservoir is formed during this time^16,17^. In the eCLEAR clinical trial, infusion of the broadly neutralizing antibody 3BNC117 at the time of ART initiation accelerated the clearance of HIV-infected cells and appeared to enhance HIV-specific CD8⁺ T-cell responses, leading to a degree of sustained ART-free control^18,19^. Interestingly, this outcome was independent of pharmacological HIV latency reversal, as an arm including the latency reversing agent (LRA) romidepsin did not further improve results.

While the development of more effective LRAs is a priority, additional studies support the idea that CTLs can impact the HIV reservoir in lieu of these. In cohorts of ART-treated individuals, HIV-specific T-cell responses are maintained in association with residual HIV expression, suggesting some ongoing recognition of infected cells^20–23^. Gradual decays of intact - but not defective - HIV proviruses over years of ART further suggest sustained immune pressure^24–26^. This selects for proviruses with an intact open reading frame for the HIV-Nef protein, which downregulates major histocompatibility class I (MHC-I)^27^ to help evade CTL immunity. In elite controllers, immune selection of the HIV reservoir is particularly pronounced and appears to take two forms: (i) selection of proviruses in poorly transcribed regions of the genome (thought to be deeply latent); and (ii) infected cells that actively transcribe HIV but possess features suggesting they can resist elimination by HIV-specific CTLs^28,29^. These observations provide a rationale to assess strategies that either counteract Nef function or target cell-intrinsic mechanisms of CTL resistance to accelerate immune-mediated reservoir decay.

One mechanism by which HIV-infected cells can resist elimination by CTLs is through overexpression of the pro-survival factor BCL-2, which is enriched in the HIV reservoir^30,31^. The BCL-2 inhibitor venetoclax enhances CTL-mediated reservoir elimination ex vivo^32,33^ and is currently being tested in a clinical trial (NCT05668026). These observations parallel findings in cancer immunotherapy, where BCL-2 drives cancer resistance to CTLs^34,35^, and venetoclax is used to treat a number of hematologic malignancies^36^.

In the current study, we identify overexpression of enhancer of zeste homolog 2 (EZH2) as a second mechanism by which HIV-infected cells can resist CTL-mediated elimination. The *EZH2* gene encodes for the catalytic subunit of the polycomb repressive complex 2 (PRC2)^37^, a histone methyltransferase that catalyzes tri-methylation of lysine 27 on histone H3 (H3K27me3). This leads to transcriptional repression, including of genes involved in MHC-I antigen presentation^38^. *EZH2* is one of the genes most frequently mutated in human lymphomas, where it is associated with lower MHC-I and MHC-II expression^39^. The EZH2 inhibitor tazemetostat, FDA approved for relapsed or refractory follicular lymphoma^40^, restores MHC-I and MHC-II expression on *EZH2*-mutant diffuse large B-cell lymphoma cells, enhancing immune surveillance of tumors^41^.

We show that *EZH2* is overexpressed in HIV-infected CD4^+^ T-cells that survive CTL co-culture. Inhibiting EZH2 by tazemetostat increases MHC-I expression on HIV-infected CD4^+^ T-cells, enhances their elimination by CTLs, and augments suppression of viral replication in vitro, in part by counterbalancing MHC-I-Nef-mediated MHC-I downregulation. In a participant-derived xenograft (PDX) mouse model, tazemetostat increases MHC-I expression enabling CTLs to reduce viral reservoir seeding and resulting in sustained skewing of CD8^+^ T-cells to less differentiated and less exhausted phenotypes. Our findings provide a rationale and mechanistic insights supporting the clinical evaluation of tazemetostat in people living with HIV.

## Results

### Overexpression of *EZH2* in cells that survive CTL exposure

We previously identified BCL-2 overexpression as a mechanism of CTL resistance by studying peptide-pulsed primary CD4⁺ T-cells that survived CTL exposure^42^. Here we sought to uncover novel mechanisms by profiling bona fide HIV-infected CD4⁺ T-cells that withstand rigorous CTL selection. CD4^+^ T-cells isolated from two people with HIV (PWH) (Supplementary Table 1) were differentiated into T_CM_ cells, and then either left uninfected or infected with the HIV isolate JR-CSF (HIV_JRCSF)_. Infected cells were either cultured with or without an autologous HIV-specific CD8^+^ CTL clone. Specifically, cells from donor OM5220 were co-cultured with a clone targeting the HIV-Env epitope ‘RLRDLLLIVTR’ (RR11), and cells from donor OM5267 were co-cultured with a clone targeting HIV-Gag ‘IRLRPGGKK’ (IK9). After overnight coculture, CD4^+^ T-cells from both conditions were sorted by flow cytometry based on infection status, determined by expression of HIV-Gag. Sorted infected (Gag^+^) and uninfected (Gag^-^) populations were subjected to transcriptional profiling via RNA-Seq (Fig. 1a).

**Fig. 1:**
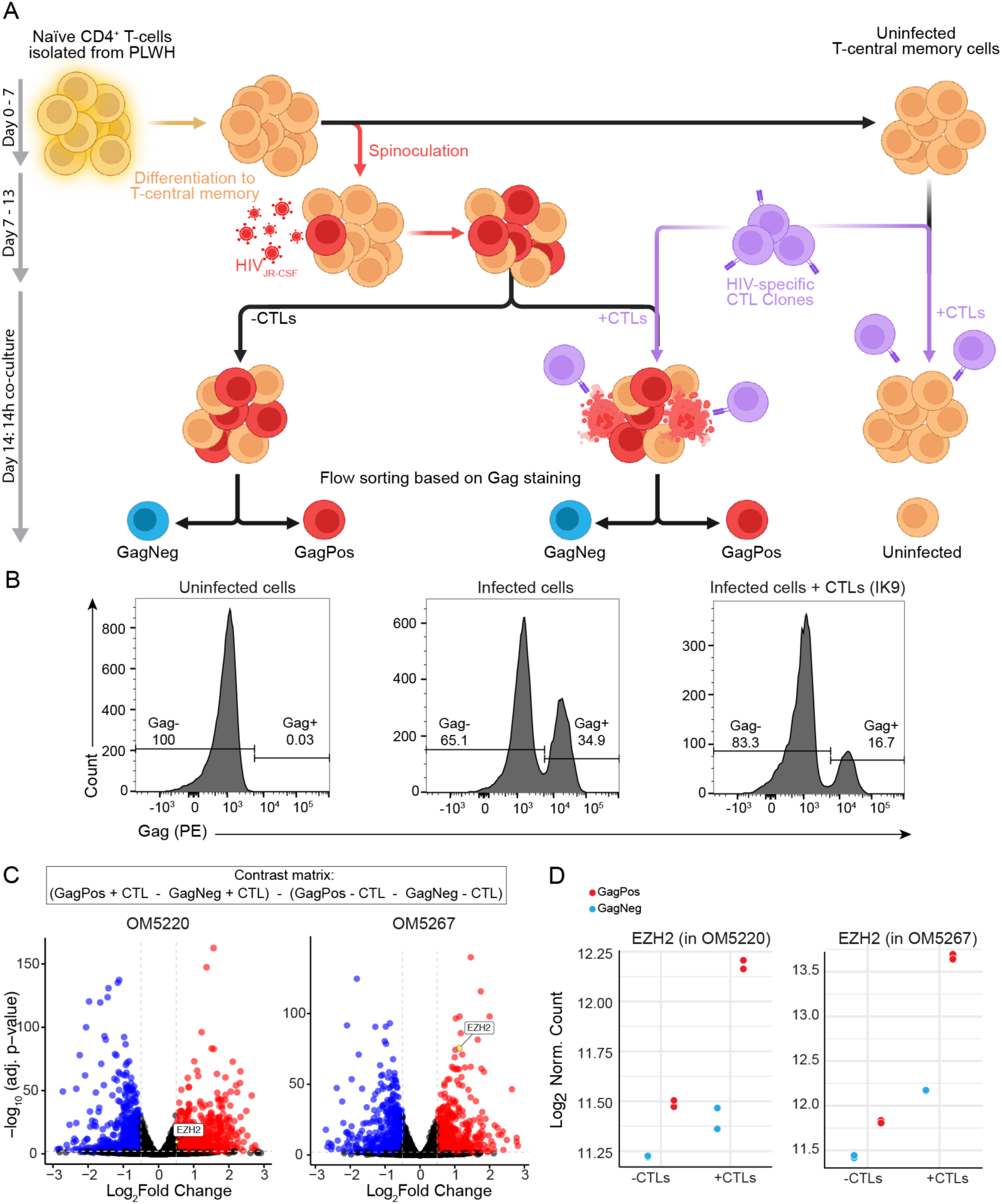
Transcriptional profiling of HIV-infected CD4^+^ T-cells that survive co-culture with autologous HIV-specific CTLs reveals *EZH2* as a candidate regulator of CTL resistance. **a)** Schematic of in vitro infected cell elimination assay and flow sorting, with timeline, prior to transcriptional profiling. **b**) Flow cytometry data gated on viable CD3^+^ lymphocytes showing the gating strategy used for flow sorting of infected (i.e., Gag^+^ or “GagPos”) cells (donor OM5267, with or without CTL clone IK9, is shown). Results show a 47% reduction in GagPos cells in +CTL condition (right panel) versus no CTL (middle panel). Refer to gating strategy in Supplementary Fig. 1. **c**) RNA-seq profiling results from infected cell elimination assay. To isolate the transcriptional signature of CTL-resistant cells and remove the impact of HIV infection itself, we applied a contrast matrix: +CTL-conditions (GagPos - GagNeg) - no-CTL conditions (GagPos - GagNeg). Shown are the resulting volcano plots for two donors (each with technical duplicates). Upregulated and downregulated differentially expressed genes (DEGs) are shown in red and blue, respectively (threshold of Benjamini-Hochberg multiple testing correction of P < 0.05). *EZH2* gene is highlighted in yellow and circled in black. **d**) Quantification of *EZH2* RNA-seq reads across all four conditions for the two donors, each in technical duplicates.

Co-culture with CTLs drove substantial reductions in frequencies of infected cells (Fig. 1b). However, two major factors complicated the identification of gene expression patterns associated with CTL resistance in this experimental system. First, cytokines produced by activated CTLs, such as interferon-gamma (IFN-γ), alter the transcriptional profiles of CD4⁺ T-cells. This makes direct comparisons between infected cells in the absence of CTLs and those that survive CTL exposure insufficient for isolating CTL resistance genes. Second, HIV infection itself induces transcriptional changes, rendering comparisons between infected and uninfected cells within the CTL-exposed condition inadequate. To address these confounders, we employed a contrast matrix (Fig. 1c) that subtracting the expression differences observed in the absence of CTLs from those observed in their presence, allowing for the identification of genes specifically associated with CTL resistance. Amongst these genes, *EZH2* stood out as a gene that was significantly overexpressed in infected cells that survived exposure to both CTL clones (adjusted P < 0.05, Benjamini-Hochberg correction) and for which FDA approved inhibitors are available^40^. Further analysis of the four parameters within the contrast matrix revealed that *EZH2* expression was only modestly impacted by HIV infection alone or by exposure to the CTL milieu alone (Fig. 1d). This expression pattern suggests that high levels of *EZH2* may specifically contribute to the survival of infected cells under CTL pressure.

### Tazemetostat increases surface expression of MHC-I on CD4^+^ T-cells

In certain cancers, *EZH2* overexpression enables immunoevasion by suppressing MHC-I expression^41,43^. We therefore investigated whether inhibiting EZH2 with the FDA approved EZH2 inhibitor tazemetostat would increase MHC-I expression on primary CD4⁺ T-cells. Tazemetostat substantially reduced H3K27me3 levels - a biomarker for EZH2 activity^44^ - in CD4⁺ T-cells (Extended Data Fig. 1a,b). We also noted impairment in CD4⁺ T-cell proliferation after 5 days of incubation with 5 μM of tazemetostat (Extended Data Fig. 1c). In-line with our hypothesis, we observed significant increases in surface MHC-I from day 3 of treatment, which intensified by day 5. Specifically, tazemetostat induced a 1.4-fold increase in MHC-I surface levels, primarily driven by upregulation of the HLA-B locus (Extended Data Fig. 1d).

### Tazemetostat enhances CTL-mediated elimination of infected CD4^+^ T-cells in vitro

Given the observed upregulation of surface MHC-I upon tazemetostat treatment, we investigated whether tazemetostat would increase the susceptibilities of HIV-infected CD4^+^ T-cells to elimination by CTLs. CD4^+^ T-cells from a PWH (participant OM5220) were infected with HIV_JRCSF,_ and then cultured in the presence or absence of the autologous HIV-specific CTL clone RR11 (Fig. 2a). Without tazemetostat, approximately 70% of target cells were eliminated at the highest effector-to-target ratio (E:T). Tazemetostat significantly enhanced the elimination of infected cells over 24 hours to approximately 90% (Fig. 2b,e). This increased susceptibility of HIV-infected cells to elimination was associated with increased MHC-I expression on CD4^+^ T-cells (Fig. 2c,d), which was observed in both infected (Gag^+^CD4^-^) and uninfected (Gag^-^ CD4^+^) T-cells (Fig. 2c). However, since MHC-I is downregulated on HIV-infected cells^45^, the net effect was to restore MHC-I levels on infected cells to more physiological levels (Fig. 2c). These findings indicate that EZH2 inhibition by tazemetostat enhances killing of HIV-infected CD4^+^ T-cells by HIV-specific CTL clones in short-term elimination assays.

**Fig. 2:**
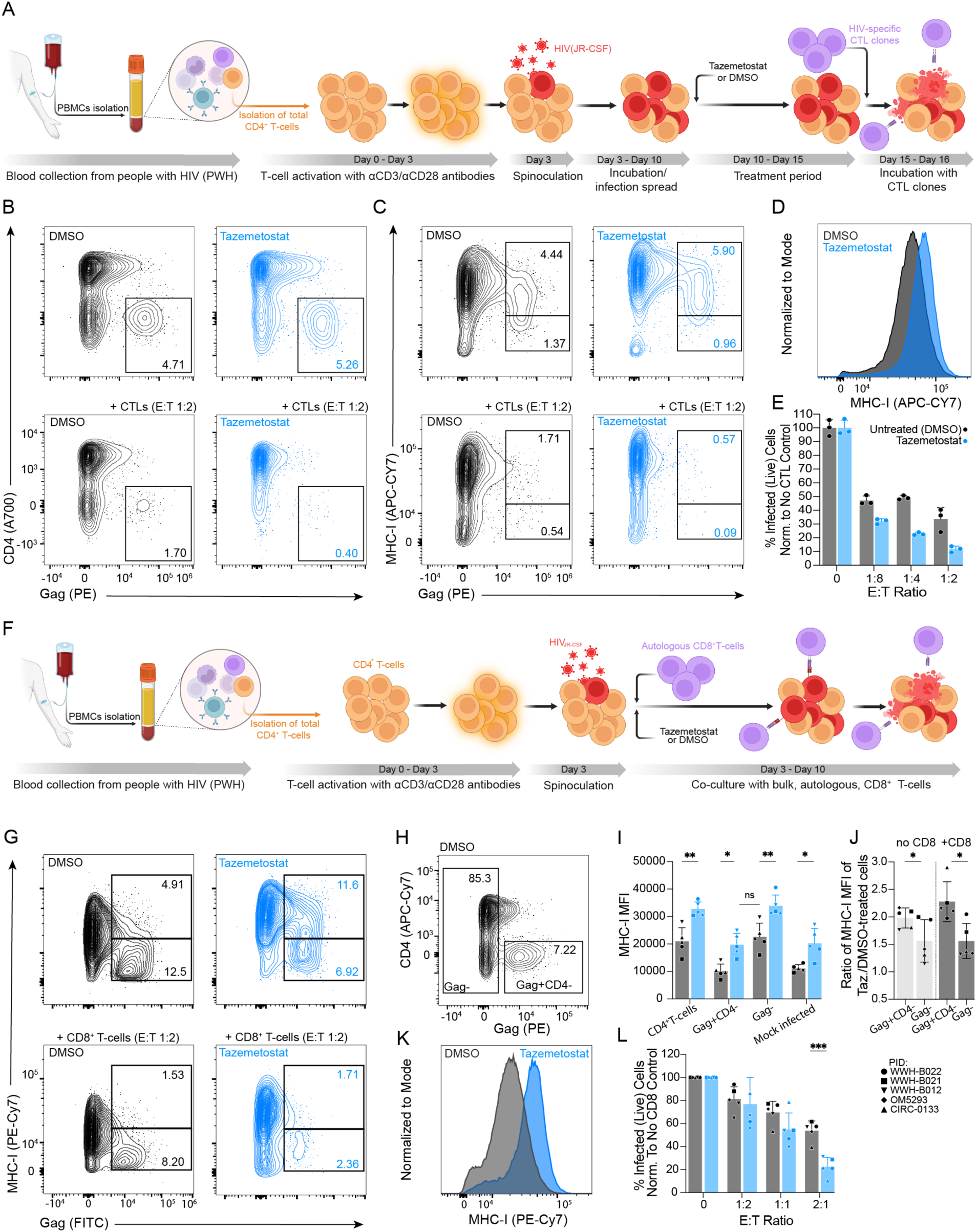
The EZH2 inhibitor tazemetostat increases surface MHC-I levels and enhances CTL-mediated elimination of HIV-infected cells in vitro. **a**) Schematic of in vitro infected cell elimination assay with tazemetostat pre-treatment used to generate the data in panels b-e. **b-c**) Flow cytometry plots depicting CD4 expression levels (b) and MHC-I expression levels (c) along with Gag expression on viable CD4^+^ T-cells pre-treated with 5 μM tazemetostat (blue) or DMSO (black), alone or after co-culture with an autologous HIV-Env-RR11-specific CTL clone. **d**) Representative histogram depicting MHC-I fluorescence intensity of CD4^+^ T-cells treated with 5μM tazemetostat (blue) or DMSO (black). **e**) Percent infected (Gag^+^CD4^-^) T-cells (within the Live-cell gate), after pre-treatment with tazemetostat (blue) or DMSO (black) and co-culture with the HIV-Env-RR11-specific CTL clone at the indicated effector:target (E:T) ratios. Dots represent n=3 technical replicates of one representative experiment. **f**) Schematic of viral inhibition assay with tazemetostat treatment used to generate the data in panels g-l. **g**) Flow cytometry plots depicting MHC-I expression levels along with Gag expression on viable CD4^+^ T-cells treated with 5 μM tazemetostat (blue) or DMSO (black), alone or after 7 days of co-culture with bulk, autologous, CD8^+^ T-cells. Refer to gating strategy in Supplementary Fig. 2. **h**) Flow cytometry plot showing gating strategy applied to define infected (Gag^+^CD4^-^) and uninfected (Gag^-^) populations, within the Live-cell gate. **i**) MHC-I fluorescence intensity of total, Gag^+^CD4^-^, Gag^-^ populations (as defined in panel h) from all (n=5) study participants. Also shown are results from ‘Mock’ cultures where virus was not added. Values are shape-coded based on participant ID (PID). Bars show mean ± SD. Statistical significance was determined by 2-way ANOVA with Šidák multiple comparison adjustment. **j**) Ratios of MHC-I MFI of tazemetostat-treated over DMSO-treated cells, in the presence or absence of CD8^+^ T-cells at 1:2 E:T ratio. Bars show mean and SD. Statistical significance was determined by Wilcoxon signed rank test. *P < 0.05. **k**) Representative histogram depicting median MHC-I fluorescence intensity of CD4^+^ T-cells treated with 5 μM tazemetostat (blue) or DMSO (black). **l**) Percent infected CD4^+^ T-cells (within the Live-cell gate) isolated from 5 PWH after treatment with 5 μM tazemetostat (blue) or DMSO (black) and co-culture with autologous CD8^+^ T-cells at the indicated E:T ratios. Bars show mean ± SD. Values are shape-coded based on PID. Statistical significance was determined by 2-way ANOVA with Šidák multiple comparison adjustment. *P < 0.05, **P < 0.01, ***P < 0.001.

To further evaluate the effect of tazemetostat on CTL-mediated control of HIV, we employed the viral inhibition assay (VIA) as a complementary in vitro approach. Unlike the previous elimination assays that utilized expanded HIV-specific CTL clones for rapid elimination of infected cells, the VIA uses ex vivo CD8⁺ T-cells as effectors and measures their ability to suppress HIV replication over a seven-day period. Since HIV-specific cells constitute a rare fraction of ex vivo CD8⁺ T-cells, effective viral inhibition in the VIA requires proliferation of these cells. Additionally, the VIA captures both cytopathic and non-cytopathic mechanisms of CD8⁺ T-cell–mediated suppression of HIV replication^46^. This assay is considered one of the most physiologically relevant in vitro models of CD8⁺ T-cell control of HIV replication, supported by studies demonstrating that VIA results correlate with in vivo HIV control^47,48^.

We performed VIAs using cells from five PWH (Supplementary Table 1). CD4⁺ T-cells were isolated, activated, and then infected with HIV_JRCSF_. These infected CD4⁺ T-cells were co-cultured with autologous CD8⁺ T-cells and either treated with 5 μM tazemetostat or maintained as untreated (DMSO) controls (Fig. 2f). At the end of the 7 day co-culture period, CD4⁺ T-cells from tazemetostat-treated cultures exhibited significantly elevated levels of surface MHC-I compared to the control condition (Fig. 2g,i,k), effectively restoring MHC-I expression on tazemetostat-treated infected cells (Gag⁺ CD4⁻) to levels similar to those of untreated uninfected cells (Fig. 2i). Notably, the fold increase in MHC-I expression upon tazemetostat treatment was greater in infected (Gag⁺) than in uninfected (Gag⁻) T-cells (Fig. 2h,j).

These increases in surface MHC-I were associated with a marked enhancement in CD8⁺ T-cell– mediated suppression of viral replication in tazemetostat-treated cultures. This manifested as greater reductions in frequencies of infected cells at higher E:T ratios (Fig. 2l). At the highest E:T ratio of 2:1, CD8⁺ T-cells reduced the percentage of infected cells by a mean of 77.5% in the tazemetostat-treated condition, compared to 46.3% in the control condition. Thus, in an assay that simulates the interplay between CD8⁺ T-cells and HIV-infected CD4⁺ T-cells, tazemetostat enhanced the capacity of ex vivo CD8⁺ T-cells to suppress HIV replication.

### Tazemetostat mitigates Nef-induced downregulation of MHC-I

The above findings suggest that tazemetostat enhances immune surveillance of HIV-infected cells by CD8^+^ T-cells through increased MHC-I antigen presentation. This model is based on the understanding that MHC-I presentation is often suboptimal on HIV-infected cells, largely due to HIV Nef, which redirects MHC-I from the Golgi apparatus to the lysosomes for degradation^49^. A prediction of this model is that tazemetostat would have less of an impact on CTL-mediated elimination of cells infected with Nef-deleted HIV. To assess this, we first confirmed that cells infected with a Nef-deleted version of HIV ‘JR-CSF(ΔNef)’ did not exhibit reduced surface MHC-I and were eliminated more efficiently than cells infected with Nef-intact virus ‘JR-CSF(WT)’ (Fig. 3a). While treatment with tazemetostat increased MHC-I levels on both JR-CSF(WT) and JR-CSF(ΔNef)-infected cells, the net effects partially restored these to physiological levels for the former versus driving supraphysiological levels for the latter (Fig. 3b,c). Correspondingly, upon overnight co-culture with an HIV-specific CTL clone, tazemetostat exhibited pronounced enhancement of elimination of cells infected with JR-CSF(WT) alongside little impact on the elimination of cells infected with JR-CSF(ΔNef) (Fig. 3d). Thus, tazemetostat acts - at least in part - by counteracting Nef-mediated immunoevasion.

**Fig. 3:**
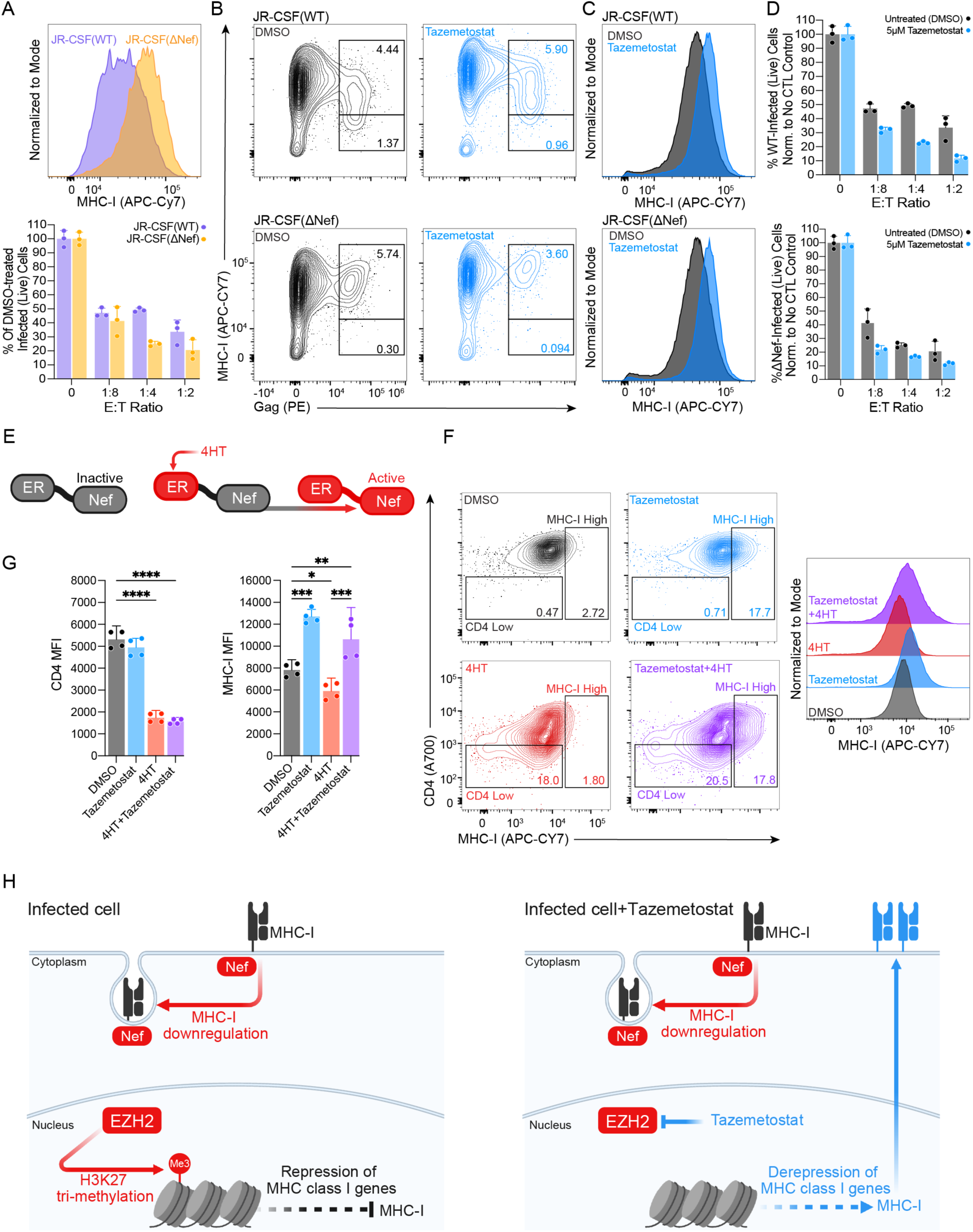
Tazemetostat counteracts Nef-mediated downregulation of MHC-I. **a**) Comparison of surface MHC-I and susceptibility to CTL between WT and ΔNef virus. Top: MHC-I fluorescence intensity of untreated CD4^+^ T-cells infected with JR-CSF(WT) (purple) or JR-CSF(ΔNef) (yellow); bottom: infected cell elimination assay shows greater reductions in cells infected with ΔNef virus. The percent infected (Gag^+^CD4^-^) T-cells (within Live-cell gate), infected with either JR-CSF(WT) or JR-CSF(ΔNef) and exposed to the HIV-Env-RR11-specific CTL clone, are shown at the indicated E:T ratios. Dots represent n=3 technical replicates of one representative experiment. **b-d**) Impact of tazemetostat on surface MHC-I and susceptibility to CTL. **b**) Flow cytometry plots depicting MHC-I levels on JR-CSF-infected (top) or JR-CSF(ΔNef)-infected (bottom) CD4^+^ T-cells treated with tazemetostat (blue) or DMSO (black). **c**) Histograms of the data shown in b, including both Gag^+^ and Gag^-^ cells. **d**) Top: Percent infected (Gag^+^CD4^-^) T-cells infected with JR-CSF(WT) (within Live-cell gate), after treatment with tazemetostat (blue) or DMSO (black) and exposure to the HIV-Env-RR11- specific CTL clone at the indicated E:T ratios; Bottom: the same experiment but with CD4^+^ T-cells infected with JR-CSF(ΔNef). Dots represent n=3 technical replicates of one representative experiment. Bars show mean ± SD. **e-f**) Isolating interplay between Nef-mediated MHC-I downregulation and tazemetostat-mediated upregulation. **e**) Schematic representation of 4HT-mediated Nef activity regulation in Sup-T1 cells. **f**) Representative flow cytometry plots of Sup-T1 cells treated as: untreated (black), tazemetostat (blue), 4HT (red), 4HT + tazemetostat (purple), and representative histograms depicting comparison of MHC-I fluorescence intensities. **g**) Comparisons of CD4 (left) and MHC-I (right) fluorescence intensity of Sup-T1 cells treated as indicated. Dots represent n=4 independent experiments. Bars show mean and SD. Statistical significance determined by ordinary 1-way ANOVA with Šidák multiple comparison adjustment. *P < 0.05, **P < 0.01, ***P < 0.001, ****P < 0.0001. **h**) Model of net effects of EZH2 and Nef on MHC-I surface expression and susceptibility to CTL. EZH2 inhibition counterbalances Nef-mediated MHC-I downregulation, sensitizing WT-virus-infected cells to CTL.

To further investigate tazemetostat in relation to Nef, we employed an uninfected Sup-T1 T-cell line that expresses a chimeric Nef-estrogen receptor (Nef-ER) hormone-binding domain protein^50^. In these cells, Nef-ER is constitutively expressed but is kept inactive until the membrane-permeable drug 4-hydroxytamoxifen (4HT) is added (Fig. 3e). In response to 48 hours incubation with 1 μM 4HT, Nef becomes activated, resulting in a 2.5-fold decrease of CD4 and MHC-I levels (Fig. 3f,g). Treatment with 5 μM tazemetostat increased surface MHC-I levels in both the absence and presence of activated Nef-ER. Notably, in cells treated with both 4HT and tazemetostat, surface MHC-I expression was restored to or exceeded physiological levels observed in untreated cells (Fig. 3f,g). This suggests that tazemetostat can counteract Nef-induced MHC-I downregulation by enhancing basal MHC-I expression. In contrast, tazemetostat had no effect on Nef-mediated downregulation of CD4 (Fig. 3f,g). These findings indicate that tazemetostat does not inhibit Nef itself but enhances MHC-I expression to levels that overcome Nef-mediated suppression (Fig. 3h). Collectively, these results demonstrate that tazemetostat can rescue MHC-I surface expression even in the presence of active Nef, potentially improving the presentation of viral antigens to CTLs.

While upregulation of MHC-I by tazemetostat sensitizes HIV-infected CD4^+^ T-cells to killing by CTLs, it could potentially impair natural killer (NK) cell responses - given that MHC-I is a ligand for inhibitory NK receptors^51^. We assessed this using in vitro NK cell elimination assays, where primary CD4^+^ T-cells isolated from 3 PWH were infected with HIV_JRCSF,_ treated or not with tazemetostat, and then cocultured with autologous NK cells. Contrary to the above hypothesis, we observed that tazemetostat treatment sensitized infected cells to improved elimination by NK cells, despite upregulation of surface MHC-I (Extended Data Fig. 2). We did not explore the underlying mechanism in this study, but one possibility could involve upregulation of NK activating ligands on CD4^+^ T-cells upon inhibition of EZH2. This would mirror observations in cancer, where EZH2 inhibition induces expression of the NK activating NKG2D ligands on hepatocellular carcinoma cells, priming these for NK-mediated elimination^52^.

### Tazemetostat Increases MHC-I and Enhances Control of HIV Viremia in vivo

We developed the HIV ‘participant-derived xenograft’ (HIV-PDX) mouse model to enable the relatively long-term in vivo evaluation of the antiviral activity of CD8^+^ T-cells taken directly from PWH^53^. We designed an initial experiment to study the impact of tazemetostat on pre-ART HIV viral load in HIV-PDX mice, following engraftment with memory CD4^+^ T-cells from a PWH (participant CIRC-0133). This individual had been on ART for 14 years at the time of sampling, and prior to this had been progressing towards AIDS (nadir CD4^+^ T-cell count 310 cells/mm^3^) (Supplementary Table 1). CIRC-0133 had HIV-specific T-cell responses that were readily detectable ex vivo by IFN-γ ELISPOT targeting HIV-Gag (440 spot forming units SFU/10^6^ PBMCs) and HIV-Pol (115 SFU/10^6^ PBMCs) (Extended Data Fig. 3). These exceeded his responses to CMV-pp65 and were in the mid to upper range of what we have previously observed in cohorts of ART-treated donors^20,21^. Mice were engrafted with memory CD4^+^ T-cells from CIRC-0133 PBMCs. 5 weeks later, mice were infected with CCR5-tropic HIV_JRCSF_ and divided into groups to receive either buffer alone (Hanks′ Balanced Salt Solution, HBSS) or autologous CD8^+^ T-cells. Each group was subdivided to receive either 500 mg/kg tazemetostat or vehicle twice a day for 27 days - overlapping for 6 days with the initiation of ART (Fig. 4a). CD8^+^ T-cells drove a sustained reduction in HIV viral load (Fig. 4b) and protected against CD4^+^ T-cell loss (Fig. 4d). Unexpectedly, tazemetostat alone was sufficient to reduce viral load by a mean of 1.75-log before ART initiation (Fig. 4c). However, the combination of CD8^+^ T-cells and tazemetostat exhibited the greatest impact, reducing viral load by a mean of 2.45-log pre-ART and facilitating undetectable viral loads in 5/8 mice by week 2 of ART treatment (Fig. 4b). The enhanced virologic control observed in tazemetostat-treated mice was associated with significant elevations in MHC-I on the surfaces of CD4^+^ T-cells within 1 week of treatment and persisting for at least 1 week after treatment was discontinued (Fig. 4e, top). MHC-I upregulation was especially pronounced in CD3^+^CD8^-^ CD4^-^ human cells, the population that contains productively HIV-infected cells (Fig. 4e, bottom).

**Fig. 4:**
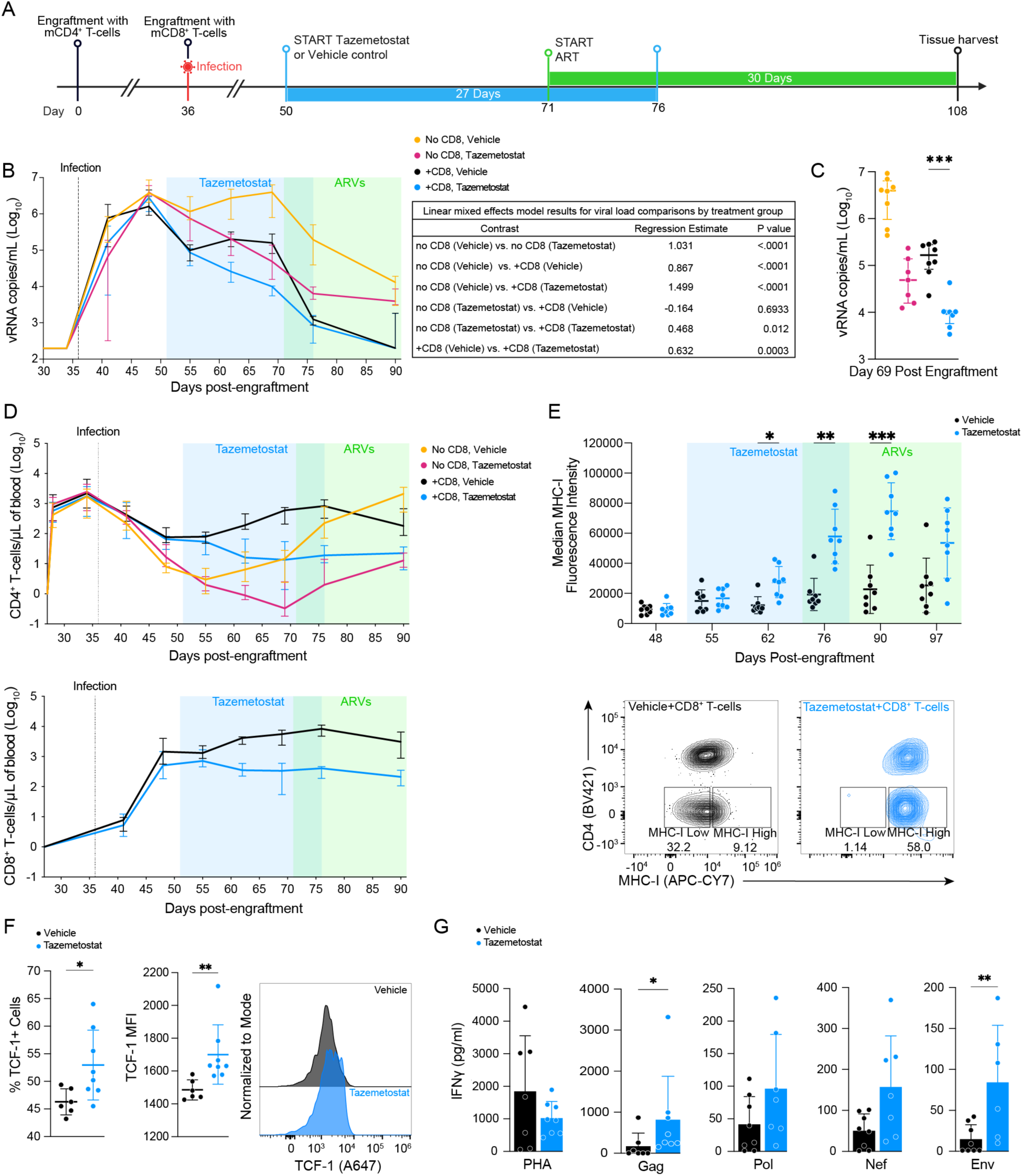
Tazemetostat enhances CD8^+^ T-cell mediated control of HIV viremia in a humanized mouse model of HIV infection. **a**) Schematic showing timeline of treatment phases in the participant-derived xenograft (PDX) humanized mouse model of HIV infection, including time of engraftment with human memory CD4^+^ T-cells, time of infection with HIVJRCSF, treatment with tazemetostat, treatment with ART, and tissue harvest. **b**) Left: Longitudinal quantification of HIV-plasma viral load in HIV-PDX mice. Shown are median with interquartile range of: vehicle-treated mice engrafted with CD4^+^ T-cells only (“no CD8” mice, yellow), tazemetostat-treated mice engrafted with CD4^+^ T-cells only (“no CD8” mice, magenta), vehicle-treated mice engrafted with CD4^+^ and CD8^+^ T-cells (“+CD8” mice, black), and tazemetostat-treated mice engrafted with CD4^+^ and CD8^+^ T-cells (“+CD8” mice, blue). Blue and green rectangles in b and c mark the timeframe of tazemetostat administration (day 50 - 76) and ARVs administration (day 71 - 107), respectively (Extended Data Fig. 4 shows viral loads of single mice). Right: Table reporting statistical comparison between groups. Linear mixed effects model includes time from infection to ART initiation (Day 36 to Day 71). Positive regression estimates represent higher viral loads in the reference group, negative estimates represent lower viral loads. P values adjusted by Tukey’s method for comparing a family of 4. **c**) HIV-plasma viral loads at day 69 post-engraftment. Each dot represents a single mouse. Shown are median with interquartile range. Values are color-coded as in b. Statistical significance was determined by unpaired two-tailed Mann Whitney U test. ***P < 0.001. **d**) Longitudinal quantification of CD4^+^ (top) and CD8^+^ (bottom) T-cell counts in HIV-PDX mice. At each time-point, median values with interquartile range are shown. Blue and green rectangles in b and c mark the timeframe of tazemetostat administration (day 50 - 76) and ARVs administration (day 71 - 107), respectively. **e**) Top: Longitudinal changes in MHC-I MFI (geomean) in CD4^+^ T-cells in +CD8 mice. Each dot represents a single mouse. Blue and green rectangles in b and c mark the timeframe of tazemetostat administration (day 50 - 76) and ARVs administration (day 71 - 107). Bars show mean and SD. Statistical significance was determined by 2-way ANOVA with Šidák multiple comparison adjustment. Bottom: Representative flow cytometry plots showing differences in MHC-I levels on CD4^+^ T-cells in vehicle- and tazemetostat- treated +CD8 mice measured at day 76 post-engraftment. **f**) Characterization of T-cell factor 1 (TCF-1) expression in CD8^+^ T-cells isolated from spleen of tazemetostat-treated (blue) or vehicle-treated (black) mice at day 107 post-engraftment. Left: %TCF-1^+^ CD8^+^ T-cells measured by flow cytometry. Center: TCF-1 MFI (geomean). Right: Representative histogram depicting TCF-1 MFI in samples from treated (tazemetostat, blue) versus untreated (vehicle, black) mice. **g**) Quantification of IFNγ released in culture supernatant of tazemetostat-treated (blue) or vehicle-treated (black) mice spleen following ex vivo incubation with either 2 μg/ml phytohemagglutinin-L (PHA) or one of the indicated HIV-peptide pools. The untreated condition (DMSO) was used for background subtraction. Each dot represents a spleen sample isolated from one single mouse. Bars show means with SD. Statistical significance was determined by unpaired two-tailed Mann Whitney U test. *P < 0.05, **P < 0.01, ***P < 0.001.

Levels of both CD4^+^ and CD8^+^ T-cells in peripheral blood were diminished in tazemetostat-treated mice in this experiment (Fig. 4d), however, CD8^+^ T-cells taken from the spleens of these mice at the time of sacrifice were skewed towards expressing T-cell factor 1 (TCF-1) (Fig. 4f), a transcription factor associated with stem-like memory T-cells. This suggests that tazemetostat treatment may preferentially expand or preserve a population of CD8^+^ T-cells with enhanced self-renewal and long-term persistence potential, which could contribute to sustained immune responses despite overall reduced circulating T-cell counts^54^. We assessed magnitudes of HIV-specific T-cell responses in splenocytes by measuring IFN-γ release following in vitro peptide stimulation. Gag-specific T-cell responses were dominant, as in pre-infusion samples, and were significantly increased in tazemetostat-treated mice. Env-specific T-cell responses were also significantly elevated in tazemetostat-treated mice, with a parallel trend for Nef and Pol. We did not observe a general increase in IFN-γ release following stimulation with the mitogen phytohemagglutinin-L (PHA) (Fig. 4g).

### Tazemetostat enables CD8-mediated reductions in HIV reservoir seeding

We extended our HIV-PDX experiments to a second donor ‘OM5220’ who had initiated ART during acute infection (Supplementary Table 1). This individual had lower ex vivo magnitudes of IFN-γ producing HIV-specific T-cell responses than CIRC-0133 with only a modest Gag-specific response of 105 SFU/10^6^ PBMCs (Extended Data Fig. 3). Following 6 weeks of memory CD4^+^ T-cell engraftment, mice were infected with HIV_JRCSF,_ and co-engrafted or not with autologous CD8^+^ T-cells. Subgroups were treated with either 500 mg/kg BID of tazemetostat or with vehicle control (Fig. 5a). The design was similar to the previous HIV-PDX experiment but including the addition of an ART interruption phase for 7/10 mice per group. The remaining 3/10 mice were sacrificed contemporaneously with ART interruption to quantify HIV reservoirs. Consistent with the previous experiment, we observed that tazemetostat alone was sufficient to reduce pre-ART HIV viral loads by an average of 1.1-log at two weeks of treatment, but with no apparent impact upon timing or levels of viral rebound upon ART interruption (Fig. 5b). CD8^+^ T-cells alone exhibited remarkable antiviral activity, intercepting early infection events to achieve an average of 0.9-log reduction in viremia 2 weeks post-infection and undetectable viremia in 6/8 mice 3 weeks later (Fig. 5b, Extended Data Fig. 5). This was accompanied by robust protection against CD4^+^ T-cell depletion (Extended Data Fig. 6; gating strategy in Supplementary Fig. 3). Following ART interruption, viral rebound was both delayed and substantially reduced in magnitude in the mice that received CD8^+^ T-cells versus the mice that did not receive CD8^+^ T-cells (Fig. 5b,c) Strikingly, all mice in the +CD8 group achieved undetectable viral loads by the end of the study, at day 192. Thus, CD8^+^ T-cells from this donor who initiated ART during acute infection exhibited exceptional control of HIV viremia in the HIV-PDX mouse model.

**Fig. 5:**
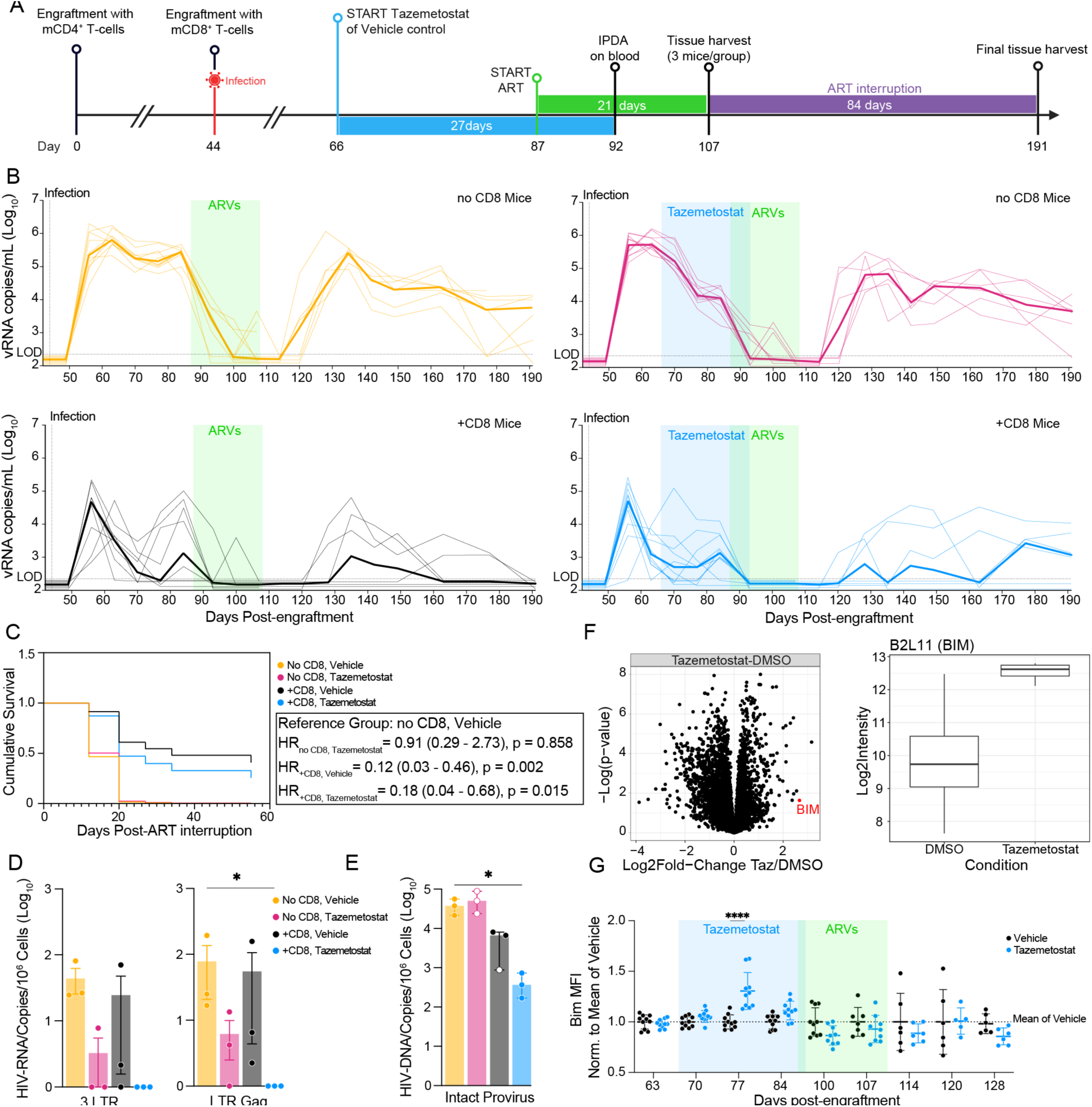
Tazemetostat in combination with CD8^+^ T-cells limits reservoir formation in the HIV-PDX model. **a**) Schematic showing timeline of treatment phases in the HIV-PDX model, including time of engraftment, time of infection with HIVJRCSF, treatment with tazemetostat, treatment with ARVs, ART interruption and tissue harvest. **b**) Longitudinal quantification of HIV plasma viral load in vehicle-treated mice engrafted with CD4^+^ T-cells only (“no CD8” mice, yellow), tazemetostat-treated mice engrafted with CD4^+^ T-cells only (“no CD8” mice, magenta), vehicle-treated mice engrafted with CD4^+^ and CD8^+^ T-cells (“+CD8” mice, black), and tazemetostat-treated mice engrafted with CD4^+^ and CD8^+^ T-cells (“+CD8” mice, blue). Thin lines represent viral load of single mice, thick lines represent median values of mice of the same group. Blue and green rectangles mark the timeframe of tazemetostat administration (day 66 - 92) and ARVs administration (day 87 - 107), respectively. **c**) Cox proportional hazards regression analysis to estimate the hazard ratio (HR) for viral rebound (>200 copies/mL), comparing to “no CD8, vehicle” as the reference group. d) Quantification of cell-associated HIV-RNA levels in spleen, measured at day 107 post-engraftment by ddPCR with primers targeting the 3’ LTR (left) or 5’ unspliced RNA (right, denoted “LTR Gag”). Bars show means with SEM. Statistical significance was determined by unpaired two-tailed Mann Whitney U test. **e**) Quantification of intact HIV DNA levels by IPDA measured at day 107 post-engraftment. Bars show means with Median and interquartile range. **f**) Left: Volcano plot displaying log2 fold changes in protein abundance between tazemetostat-treated and DMSO conditions (tazemetostat/DMSO) from the same samples used in the viral inhibition assay (Fig. 2). Each point represents a protein; proteins significantly altered by tazemetostat treatment are indicated by the log2 fold change and -log10(p-value). The BIM protein is highlighted in red. Right: Box plot showing the log2 intensity of B2L11 (BIM) protein levels in DMSO and tazemetostat-treated conditions (additional mass spectrometry data in Extended Data Fig. 8) **g**) Longitudinal changes in BIM MFI (geomean) in the CD4^+^ T-cell population of +CD8 mice that were either treated with tazemetostat (blue) or vehicle (black). MFI values were normalized to the mean of vehicle-treated mice. Each dot represents a single mouse. Blue and green rectangles mark the timeframe of tazemetostat administration (day 66 - 92) and ARVs administration (day 87 - 107), respectively. Statistical significance was determined by 2-way ANOVA with Bonferroni multiple comparison adjustment. *P < 0.05, **P < 0.01, ***P < 0.001, ****P < 0.0001.

In contrast to the previous experiment (Fig. 4) we did not observe a further impact on HIV viral load in mice that received tazemetostat in addition to CD8^+^ T-cells, despite confirming potent in vivo reductions in H3K27me3 levels and upregulation of MHC-I in CD4^+^ T-cells (Extended Data Fig. 7). We reasoned that the exceptional performance of CD8^+^ T-cells in this experiment may have left little room for further improvement in viral loads with tazemetostat, but that enhanced killing of HIV-infected cells may yet be reflected in the seeding of the HIV reservoir. To assess this, we studied splenocytes from the 3 mice per group sacrificed on day 107, just prior to ART interruption. We first quantified levels of cell-associated polyadenylated HIV RNA using two sets of primers and probes targeting either the 5’ or 3’ region of the HIV genome. The combination of CD8^+^ T-cells with tazemetostat was unique in achieving a significant reduction in 5’ (LTR gag) HIV mRNA (p = 0.04) with a parallel trend for 3’ (3LTR) HIV mRNA (p = 0.08), manifesting as a lack of detectable HIV mRNA in mice from this group (Fig. 5d). We then applied the intact proviral DNA assay (IPDA) to quantify putatively genome intact HIV proviruses^55^. As with HIV mRNA, the group that received both CD8^+^ T-cells and tazemetostat was unique in achieving significantly lower levels of intact HIV proviruses than the CD4 only control group (Fig. 5e). This may be a result of the observed upregulation of MHC-I on CD4^+^ T-cells. However, we were led to explore another putative mechanism by proteomics analysis performed on CD4^+^ T-cells from the viral inhibition assays featured in Fig. 2, which revealed over-expression of the proapoptotic BCL-2 family member BIM following tazemetostat treatment (Fig. 5f) (proteomics analysis in Extended Data Fig. 8). In the current mouse experiment, we observed a marked increase in BIM in the CD4^+^ T-cells of the tazemetostat-treated groups, both with (Fig. 5g) and without CD8^+^ T-cells (data not shown). Since over-expression of pro-survival BCL-2 can confer CTL resistance in the HIV reservoir^30^, the induction of BIM expression by tazemetostat may therefore serve to counterbalance this and sensitize infected cells to clearance.

### Persistent reduction in PD-1 and skewing toward a T_SCM_ phenotype in CD8^+^ T-cells from tazemetostat-treated PDX mice

The increased TCF-1 expression observed at the conclusion of the previous mouse experiment (using CIRC-0133) was notable, given that TCF-1 expression in HIV-specific CD8^+^ T-cells has been closely linked with immune control of both HIV and SIV^56^. TCF-1^+^ cells display stem-like properties, including robust expansion and self-renewal, which are critical for sustained immune responses to these viruses^54^. To investigate this further, we assessed TCF-1 in the second mouse experiment, and beginning on day 135, we also assessed CD45RA, CD27, CCR7, and CD95, to identify key CD8^+^ T-cell subsets: naive, stem cell memory T_SCM,_ central memory T_CM,_ effector memory T_EM,_ transitional memory T_TM,_ terminal effector T_TE_^57,58^ ^(^Fig. 6a). Mice treated with tazemetostat showed a consistent trend towards increased proportions of T_SCM_ CD8^+^ T-cells, which reached statistical significance at days 149 and 163 (Fig. 6b,c). This increase coincided with significantly higher percentages of TCF-1^+^ CD8^+^ T-cells at days 135, 142, and 163 post-engraftment (Fig. 6d), corroborating our earlier results. As tazemetostat was administered only from days 66-92, these results are indicative of a durable immune restoration in favor of more stem-like CD8^+^ T-cells. Similar to TCF-1 expression, higher frequencies T_SCM_ CD8^+^ T-cells are strongly associated with immune control and improved prognosis in untreated HIV infection^56^. Thus, beyond reducing HIV reservoirs, treatment with tazemetostat induced a long-lasting shift in CD8^+^ T-cells towards a profile associated with immune control.

**Fig. 6:**
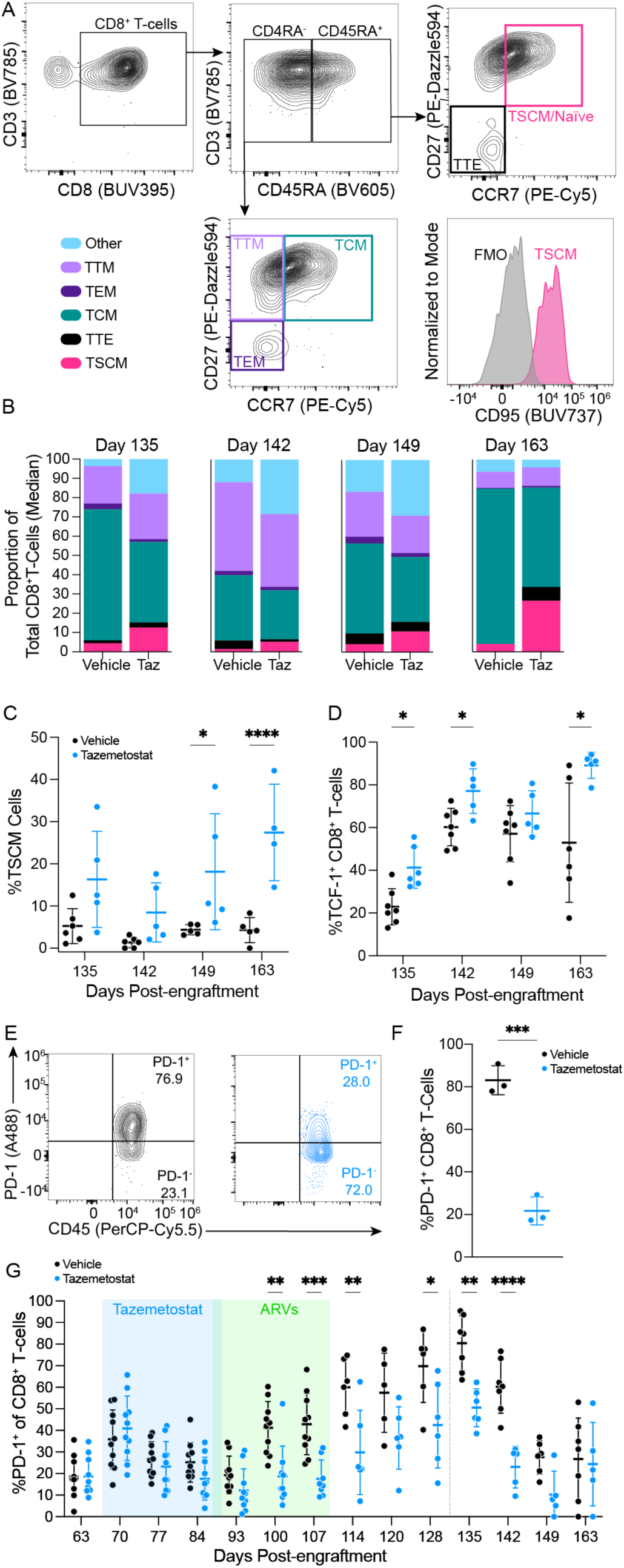
CD8^+^ T-cells in tazemetostat-treated mice are skewed to a more stem-like phenotypic profile. **a**) Flow cytometry gating strategy used to define CD8^+^ sub-populations. Gating is color-coded: T-transitional memory (TTM, light purple); T-central memory (TCM, green), T-effector memory (TEM, dark purple); T-stem cell memory (TSCM/Naïve, magenta); T-terminal effector (TTE, black). Cells were gated as shown in Supplementary Fig. 4. Histogram shows expression of CD95 in a representative sample and FMO control. **b**) Stacked bar charts reporting proportions of CD8^+^ sub-populations in tazemetostat-treated or vehicle-treated mice at the indicated time-points. **c**) Percent TSCM cells in tazemetostat-treated (blue) or vehicle-treated (black) mice at the indicated time-points. Each dot represents a single mouse. Bars show mean ± SD. Statistical significance was determined by 2-way ANOVA with Bonferroni multiple comparison adjustment. **d**) Percent PD-1^+^ CD8^+^T-cells in spleen of tazemetostat-treated (blue) or vehicle-treated (black) mice at day 107 post-engraftment (cells were gated as shown in Supplementary Fig. 3b). Shown are mean ± SD. Statistical significance was determined by unpaired, parametric, two-tailed t test. ***P < 0.001. **e**) Representative flow cytometry plots showing changes in %PD-1^+^ and PD-1^-^ CD8^+^T-cells in blood of tazemetostat-treated (blue) or vehicle-treated (black) mice; plots show concatenated values of n=7 mice (Vehicle) and n=6 mice (tazemetostat) at day 128 post-engraftment (gated as shown in Supplementary Fig. 3b). **f**) Percent PD-1^+^ CD8^+^T-cells in spleen of tazemetostat-treated (blue) or vehicle-treated (black) mice at day 107 post-engraftment (cells were gated as shown in Supplementary Fig. 3b). Bars show mean ± SD. Statistical significance was determined by multiple unpaired two-tailed t test, with Holm-Šidák multiple comparison adjustment. **g**) Longitudinal changes in %PD-1^+^ CD8^+^ T-cells in blood samples of +CD8 mice. Each dot represents a single mouse treated either with tazemetostat (blue) or vehicle (black) at the indicated time-points. Blue and green rectangles mark the timeframe of tazemetostat administration (day 66 - 92) and ARVs administration (day 87 - 107), respectively. Time points to the right of the dotted line were determined with the flow cytometry panel used to define CD8^+^ populations. Bars show mean and SD. Statistical significance was determined by 2-way ANOVA with multiple comparison adjustment. **P < 0.01, ***P < 0.001, ****P < 0.0001.

To further evaluate whether tazemetostat had reinvigorated CD8^+^ T-cells, we next turned our attention to PD-1, a co-inhibitory receptor that plays a central role in HIV-induced T-cell exhaustion^59–61^. Looking first at cells from the spleens of the 6 mice sacrificed on day 107 (just prior to ART interruption), PD-1 levels were found to be dramatically reduced in tazemetostat-treated mice (mean ± SD: Vehicle - 83.1 ± 6.8% PD-1^+^, tazemetostat - 21.7 ± 6.6, p < 0.001, Fig. 6f). Extending these findings to the blood, CD8^+^ T-cells from tazemetostat-treated mice exhibited significantly lower PD-1 expression starting at day 100, while mice were still suppressed on ART, and extending to day 142 (Fig. 6g). Mice were followed with this same flow cytometry panel out to day 128, where tazemetostat-treated mice continued to retain lower PD-1 expression (Fig. 6g). From day 135 to 163, we switched to a different flow cytometry panel (in order to characterize maturational phenotypes) and continued to observe reduced PD-1 expression in tazemetostat-treated mice (Fig. 6g). Thus, transient treatment with tazemetostat just prior to ART initiation resulted in durable improvements in clinically relevant phenotypes in CD8^+^ T-cells taken directly from PWH.

## Discussion

In this study, we identified naturally occurring heterogeneity in EZH2 expression as a novel determinant of the susceptibilities of HIV-infected CD4^+^ T-cells to elimination by CTL. By inhibiting EZH2 with the FDA-approved drug tazemetostat, we demonstrated enhanced CTL-mediated killing of HIV-infected cells in vitro and improved control of viral replication by CD4^+^ T-cells during longer-term co-cultures. These effects were associated with upregulation of MHC-I expression on CD4^+^ T-cells, effectively counteracting HIV Nef-mediated MHC-I downregulation - a key immune evasion strategy of the virus.

Our in vivo studies using the PDX mouse model further supported these findings. Tazemetostat treatment enhanced CD8^+^ T-cell-mediated control of pre-ART viral load in one donor and reduced HIV reservoir seeding in another donor with potent baseline CD8^+^ T-cell responses. Notably, tazemetostat-treated mice exhibited favorable alterations in CD8^+^ T-cell phenotypes, including lower PD-1 expression and a skewing towards a T_SCM_ phenotype that persisted for at least 79 days after the conclusion of treatment. These observations suggest that EZH2 inhibition not only sensitizes infected cells to immune clearance, but may also promote long-term immune surveillance by enhancing CD8^+^ T-cell function and persistence.

A direct implication of our results is to motivate studies assessing whether treatment with tazemetostat around the time of ART initiation may be sufficient to achieve beneficial virologic and immunologic outcomes in PWH, namely reduced HIV reservoir sizes and lower immune activation, as measured by PD-1. These in turn, might impact the inflammation-associated comorbidities that persist in ART-treated individuals, such as cardiovascular disease^62–65^. Decreasing HIV reservoirs and improving the T_SCM_ profiles of CD8^+^ T-cells are also key upstream goals of efforts to cure HIV infection, defined either as eradicating HIV reservoirs or empowering the immune system to control viral rebound^66^. While we did not observe an impact of tazemetostat on viral rebound in the current study, several limitations are notable here. First, the short duration of ART in our studies does not reflect a likely clinical scenario, where any interruption would most likely occur after months or years of treatment. Whereas a reduction in the size of a stable/quiescent reservoir may lead to a delay in HIV rebound, short-term treatment may leave many infected cells poised to reseed replication. Second, in relation to post-rebound control of viremia, we observed a remarkable degree of CD8-mediated control (that would be enviable in a human clinical setting) in our vehicle control condition. Although CD8^+^ T-cells in the tazemetostat-treated group exhibited skewing towards favorable phenotypic profiles, there may have been little room to improve upon the control observed in the CD8 + vehicle group. The current study substantially pushed the boundaries of the HIV-PDX model itself, which has not previously been used to demonstrate CD8-mediated control of post-ART viral rebound. A notable additional side observation is the ability of the model to reveal a remarkably effective CD8^+^ T-cell response that would not have been anticipated based on relatively weak ELISPOT responses. Future studies with tazemetostat in HIV-PDX mice could extend to greater variety of donors, including those with CD8^+^ T-cell responses that are less effective at baseline, or could alter a number of parameters to detune CD8^+^ T-cell efficacy (e.g. longer duration of pre-ART viremia, longer duration of ART treatment, engraftment with fewer CD8^+^ T-cells, and others).

While we have focused on the enhancement of MHC-I antigen presentation by tazemetostat, and also presented its ability to skew the BCL-2 family axis to favor apoptosis, the inhibition of EZH2 has a number of additional modalities of action by which it may contribute to curing HIV infection. EZH2 is known to guide effector CD8^+^ T-cell terminal differentiation and loss of multipotency by initiating and maintaining methylation of pro-memory and pro-survival genes^67^. Thus, while the skewing that we observed in tazemetostat-treated mice towards PD-1^-^ T_SCM_ CD8^+^ cells may be a result of diminished antigen exposure from reduced HIV reservoirs, it could also reflect a direct epigenetic reprogramming of CD8^+^ T-cells. Epigenetic modification by EZH2 has also been reported to play a critical role in the establishment and maintenance of HIV latency^68–71^. Thus, tazemetostat may help expose these infected cells to immune clearance. Finally, EZH2 plays a critical role in promoting cell proliferation in a number of contexts^72,73^. Our results show that tazemetostat impairs CD4^+^ T-cell proliferation in vitro. We propose that the enhanced CD4^+^ T-cell depletion and reduced HIV viral load in mice that received tazemetostat only (no CD8^+^ T-cells) may reflect impaired replenishment of CD4^+^ T-cells through reduced proliferation. While not desirable in the context of uncontrolled HIV, in the setting of CD8-mediated virologic control during ART, reducing CD4^+^ T-cell proliferation may curb the clonal expansion of HIV-infected cells that contributes to HIV persistence^74–77^. Further studies are needed to optimize efforts to leverage these additional putative benefits of EZH2 inhibition, alongside the enhanced susceptibility to CTL demonstrated in the current study.

Achieving either HIV eradication or remission is likely to require treatment with a combination of agents. Based on our findings, several potential strategies merit exploration - in either the HIV-PDX mouse, other humanized mouse models, and/or non-human primates. Combining tazemetostat with immunotherapeutic agents like the IL-15 super-agonist N-803 could enhance CD8^+^ T-cell effector functions alongside sensitizing infected cells to killing^78^. Integration with broadly neutralizing antibodies (bNAbs) may offer a multifaceted approach to reducing the reservoir, as bNAbs have shown promise in limiting reservoir seeding and enhancing CD8^+^ T-cell-mediated control^18,19^. Given the role of EZH2 in maintaining HIV latency^68–71^, combination with LRAs with synergistic modes of action could effectively expose latent reservoirs to immune clearance. Additionally, using BCL-2 inhibitors like venetoclax alongside tazemetostat could target multiple resistance mechanisms in HIV-infected CD4⁺ T-cells, as this combination has demonstrated synergistic effects in sensitizing lymphoma cells to apoptosis^79^.

In conclusion, our study provides mechanistic insights into how EZH2 overexpression enables HIV-infected cells to evade immune clearance and demonstrates the potential of tazemetostat to enhance CTL-mediated elimination of these cells. By increasing MHC-I expression and modulating key apoptotic pathways, tazemetostat addresses critical barriers to HIV reservoir reduction. The sustained improvement in CD8⁺ T-cell phenotypes further supports its potential role in cure strategies. We believe that clinical evaluation of tazemetostat in PWH is warranted to determine its efficacy and safety in this context. Integrating EZH2 inhibition into combination therapies could be a promising approach toward achieving durable viral remission or eradication, ultimately bringing us closer to a scalable and effective HIV cure.

## Methods

### Data reporting

Mouse group sizes were determined based on power calculations provided in our previous publication^53^. The experiments were not randomized, and the Investigators were not blinded to allocation during experiments and outcome assessment.

### Statistics

Statistical analyses including logarithmic transformations, t-tests, Wilcoxon tests, and two-way ANOVAs were performed using GraphPad Prism 10. Linear mixed-effects (LME) models were conducted using the “lme4” package in R^80^. B-spline functions of time with interior knots at each sampling time point were used in LME models, and terms for correlated random intercepts and random slopes were included in models. Multiple comparisons within LME models were made using the “emmeans” package in R (Lenth R, 2024. *emmeans: Estimated Marginal Means, aka Least-Squares Means*. R package version), adjusting for multiple comparisons using Tukey’s all-pair method. Statistical tests used are indicated in figure legends.

### Participants and study approval

Study participants were recruited from the Maple Leaf Medical Clinic (Toronto, Canada) or from Whitman Walker Health (Washington, D.C). Apheresis from HIV negative individuals was collected from the Gulf Coast Regional Blood Center. Ethical approval to conduct this study was obtained from University of Toronto, George Washington University, and the Weill Cornell Medicine Institutional Review Boards. All participants were adults and provided written informed consent.

### PBMC and cell subset isolation

Peripheral blood mononuclear cells (PBMCs) were isolated from leukapheresis and apheresis samples via standard Ficoll gradient separation and were used immediately or cryopreserved (- 150°C, 90% Fetal Bovine Serum + 10% DMSO). Total, memory, or naïve CD4⁺ T-cells were isolated from PBMCs using the EasySep Human CD4^+^ T-Cell Enrichment Kit, the EasySep Human Memory CD4^+^ T-Cell Enrichment Kit, or the EasySep Human naïve T-cell isolation kit (Stemcell Technologies), respectively, according to the manufacturer’s recommendations, with the exception that an initial cell concentration of 10^8^ cells/mL was used.

### Media and reagents

The following culture media were used: R10 (RPMI 1640 supplemented with 10% FBS and 1% penicillin/streptomycin, 1% HEPES buffer, 1% L-Glutamine), R10-50 (R10 supplemented with 50 Units/mL of IL-2), R10-50 + IL15 (R10-50 supplemented with 10ng/mL of IL-15).

### Preparation of HIV stocks

JR-CSF(ΔNef) was created from the JR-CSF plasmid (NIH AIDS reagent program) though a 20 amino acid deletion after serine 56, which produced a stop codon immediately after the serine and introduced multiple downstream stop codons downstream due to the frameshift. Infectious viral stocks of HIV_JRCSF_ or HIV_JRCSF_(ΔNef) were generated by transfecting full-length plasmids into HEK293T cells (ATCC, CRL-3216) using FuGENE 6 Transfection Reagent (Promega). 2 days after transfection, tissue culture media were collected, pre-cleared by centrifugation at 2,000 x g for 5 min, sterile filtered through a 0.45-μm syringe filter and concentrated using PEG-it Virus Precipitation Solution (System Biosciences). Virus was then aliquoted and stored at −80°C. Virus titration was performed on TZM-bl cells as described previously^81^.

### In vitro infected-cell elimination and isolation of survivor HIV-infected cells

Isolated CD4^+^ T-cells were differentiated to central memory (T_CM_) populations adapting a method previously described^82,83^. Briefly, isolated naïve CD4^+^ T-cells were activated using human anti-CD3/anti-CD28-coated magnetic beads (one bead for two cells, 11131D, Gibco) in the presence of human aIL-4 (1ug/10^6^, 500-P24, Peprotech), human aIL-12 (2ug/10^6^, 500-P154G, Peprotech), and Tumor Growth Factor (TGF)-b1 (10ng/10^6^, 100-21, Peprotech) to prevent cell polarization. After 3 days, cells were maintained at the concentration of 10^6^/mL in media containing 50 UI of human IL-2. At day 7, the cultured T_CM_ were infected with HIV-1_JRCSF_. One quarter of the cells were infected by spinoculation with 200 μl of virus for 10 million cells in 1mL and centrifuged at x 1500g for 2h at 37°C. Following the spinoculation, the infected cells were mixed in a flask with the remaining three quarters of cells at 2x10^6^/mL in R10-50 media. At day 10, the cells were crowded in 96-well round bottom plates using a density of 2x10^5^ in 200uL/well to enhance cell-to-cell transmission. On day 13, infected cells were then cultured alone or with autologous HIV-specific CD8^+^ CTL clones for 14 hours at the noted effector to target (E:T) ratio. Cells isolated from donor OM5220 were co-cultured with a CTL clone targeting the HIV-Env epitope ‘RLRDLLLIVTR’ (RR11), and cells isolated from donor OM5267 were co-cultured with a CTL clone targeting the HIV-Gag epitope ‘IRLRPGGKK’ (IK9). Subsequently, cells were stained for flow cytometry analysis with the following human-specific antibodies: CD4 (AF488, RPA-T4, Biolegend); CD3 (BV785, OKT3, Biolegend); CD8 (BV605, RPA-T8, Biolegend); anti-HIV-1 core antigen (RD1, KC57, Beckman Coulter). Live/Dead Aqua Fixable cell stain (Invitrogen) was used as a viability marker. Cells were fixed and permeabilized using the cytofix/cytoperm (BD) fixation/permeabilization kit. Single Gag^+^ cells were sorted on the BD FACsAria Cell Sorter with the strategy represented in Supplementary Fig. 1. BD FACSDiva Software v8.0.2 was used to acquire flow cytometry data and Flow-Jo software v10.6.2 was used to analyze flow cytometry data (Tree-Star).

### RNA-Seq sample acquisition

Total RNA was isolated from FACs-sorted PFA-fixed cells (populations as shown in Supplementary Fig. 1) using the miRNAeasy FFPE kit for microRNA extraction (Qiagen). Total RNA integrity was assessed using the Agilent Bioanalyzer, libraries were generated for samples with a RIN >8.0 with the TruSeq RNA sample Preparation kit (Illumina). cDNA libraries were sequenced with the HISeq4000 instrument.

### RNA-Seq data analysis

Raw reads were quality checked with FastQC v0.11.7 (http://www.bioinformatics.babraham.ac.uk/projects/fastqc/). Reads were aligned to the human reference genome (GRCh38.p12) with STAR v2.6.0.c^84^. Gene abundances were calculated with featureCounts v1.6.2 using composite gene models from Gencode release 28^85,86^. Differentially expressed genes were determined with DESeq2 v1.32.0 using Wald tests^87^. Corrected P values were calculated based on the Benjamini-Hochberg method to adjust for multiple testing. The list of differentially expressed genes comparing “real survivors” and “real bystanders” was determined with a cutoff of FDR (adjusted P value) of less than 0.01. Gene counts from RNA-Seq data was deposited to GitHub (https://github.com/abcwcm/Gramatica_Miller2024.git).

### In vitro and in vivo treatment with tazemetostat

In vitro: tazemetostat (EPZ-6438, Selleckchem cat #S7128) was purchased as powder, diluted in DMSO (up to 100 mg/mL), aliquoted and kept frozen until use. Whenever indicated, DMSO alone was used as negative control, up to 0.01% (vol/vol) final volume. In all in vitro experiments, fresh tazemetostat was replenished in the cell culture every other day. In vivo: tazemetostat (EPZ-6438, Glpbio,GC14062) was kept lyophilized and frozen until use. Aliquots of tazemetostat were prepared prior to each oral administration, by diluting the appropriate amount of lyophilized compound in vehicle (0.5% sodium carboxymethyl cellulose + 0.1% Tween 80) and sonicating until homogenized.

### Biomarker and MHC-I analysis on CD4^+^ T-cells

Isolated CD4^+^ T-cells from HIV negative donors were then activated for 72 hours in the presence of 1 μg/mL each anti-CD3 (eBioscience, OKT3) and anti-CD28 (eBioscience,CD28.2) antibodies in R10-50, washed and, either left untreated (DMSO control), or treated with the indicated concentration of tazememtostat for 5 days. Subsequently, cells were stained for flow cytometry analysis with the following human-specific antibodies: HLA-A (BV605, 1082C5, BD), HLA-B (BV650, YTH 76.3.rMAb, BD), HLA-E (PerCP/Cyanine5.5, 3D12, Biolegend), HLA-C (APC/Cyanine7, DT-9, Novusbio), CD4 (BV421,RPA-T4, Biolegend), HLA-A,B,C, (APC/Cyanine 7, W6/32, Novusbio), anti-Tri-Methyl-Histone H3 (Lys27) (PE-Cy7, C36B11, CST), CD3 (BV785, OKT3, Biolegend). Live/Dead Aqua Fixable cell stain (Invitrogen) was used as a viability marker. Cells were fixed and permeabilized using the Foxp3/Transcription Factor Staining Buffer Set (eBioscience). Samples were analyzed on an Attune NxT flow cytometer and Flowjo software (TreeStar).

### In vitro infected-cell elimination assay

These methods apply to the results shown in Figs. 2 & 3. CD4^+^ T-cells were isolated from participant OM5220, activated for 72 hours with 1 μg/mL anti-CD3 (OKT3, eBioscience) and anti-CD28 (CD28.2, eBioscience) antibodies in R10-50, and subsequently spinoculated for 1.5 hours at 1,200g (37°C) with either HIV_JRCSF_ or HIV_JRCSF_ (ΔNef). Cells were then incubated for 5 days with either 5μM tazemetostat or DMSO (untreated control). On the following day, CD4^+^ T-cells were then co-cultured overnight in the presence of CFSE- or CellTraceFarRed (CTFR)-labeled RR1(RLRDLLLIVTR)-specific CTL clones at the indicated E:T ratios (CFSE, CTFR sourced from Invitrogen). Subsequently, cells were stained for flow cytometry analysis with the following anti-human antibodies: CD4 (BV421, RPA-T4, Biolegend), HLA-A,B,C (PE-Cy7, W6/32, Biolegend), anti-Tri-Methyl-Histone H3 (Lys27) (PE-Cy7, C36B11, CST), CD3 (BV785, OKT3, Biolegend), anti-HIV core antigen (RD1, clone KC57, Beckman Coulter). Antibodies were added at a 1:100 dilution. Live/Dead Aqua Fixable cell stain (Invitrogen) was used as a cell viability marker. Cells were fixed and permeabilized using the Foxp3/Transcription Factor Staining Buffer Set (eBioscience). Samples were analyzed on an Attune NxT flow cytometer and Flowjo software (TreeStar).

### Viral inhibition assay

Isolated CD4^+^ T-cells were activated for 72 hours with 1 μg/mL anti-CD3 (OKT3, eBioscience) and anti-CD28 (CD28.2, eBioscience) antibodies in R10-50, and subsequently spinoculated for 1.5 hours at 1,200 x g (37°C) with HIV_JRCSF._ On the same day, autologous CD8^+^ T-cells were isolated from cryopreserved PBMCs and labeled with CTFR. Following spinoculation, CD4^+^ T-cells were plated either alone or with autologous CD8^+^ T-cells at the indicated E:T ratios and incubated at 37°C, 5% CO_2_ for 7 days. Where indicated, co-cultures of targets and effectors were incubated with either 5 μM tazemetostat or DMSO (untreated control). Co-cultures were then stained for flow cytometry analysis with the following anti-human antibodies: CD3 (BV785, OKT3, Biolgened), anti-CD4 (APC/Cyanine7, RPA-T4, Biolegend), HLA-A,B,C (PE/Cyanine7, W6/32, Biolegend), CD8 (BV605, SK1, Biolegend), and anti-HIV core antigen (RD1, KC57, Beckman Coulter). Antibodies were added at a 1:100 dilution. Live/Dead Aqua Fixable cell stain (Invitrogen) was used as a viability marker. Cells were fixed and permeabilized using the Foxp3/Transcription Factor Staining Buffer Set (eBioscience). Samples were analyzed on an Attune NxT flow cytometer and Flowjo software (TreeStar).

### MHC-I and CD4 analysis on Nef-inducible Sup-T1 cells

Nef-ER expressing Sup-T1 cells (Clone 31) were obtained through the NIH HIV Reagent Program, Division of AIDS, NIAID, NIH: Nef-ER Expressing Sup-T1 Cells (Clone 31), ARP-6453, contributed by Drs. Scott Walk, Kodi Ravichandran and David Rekosh^50^. Cells were grown in R10 media. To activate intracellular Nef, cells were treated with 1μM 4-hydroxytamoxifen (Selleckchem) for 48 hours. Subsequently, either tazemetostat or DMSO were added, and the cells were cultured for 5 days. Following treatment, cells were stained for flow cytometry analysis with the following human specific antibodies: HLA-A,B,C(PE-Cyanine7, W6/32, Biolegend), CD4 (BV421, RPA-T4, Biolegend), CD3 (BV785, OKT3, Biolegend). All antibodies were added at 1:100 dilutions. Live/Dead Aqua Fixable cell stain (Invitrogen) was used as a viability maker. Samples were analyzed on an Attune NxT flow cytometer and Flowjo software (TreeStar).

### Sample preparation for proteomics analyses

In parallel to each VIA (Fig. 2f-l), 1-2 x 10^6^ of the same CD4^+^ T-cells isolated from 5 PWH (PIDs: OM5220, OM5293, CIRC-0133, WWH-012, WWH-021, WWH-022) were set aside, plated at 1 x 10^6^ cells/ml and treated with 5 μM tazemetostat or left untreated (DMSO) for 7 days. Subsequently, cells were lysed in a buffer comprised of 8 M Urea (Sigma), 50 mM ammonium bicarbonate (Sigma), 150 mM NaCl (Sigma), and protease/phosphatase inhibitor cocktail (HALT, Thermo Scientific) prepared in Optima LC/MS grade water (Fisher Scientific). Lysates were probe-sonicated on ice three times for 1 second at 50% power, with 5 seconds of rest in between pulses. Protein content of the lysates was quantified using a micro-BCA assay (Thermo Fisher). Samples were treated with Tris-(2-carboxyethyl)phosphine to a final concentration of 4 mM and incubated at room temperature (RT) for 30 minutes. Iodoacetamide (IAA) was added to each sample to a final concentration of 10 mM, and samples were incubated in the dark at RT for 30 minutes. IAA was quenched by 10 mM dithiothreitol and incubated in the dark at RT for 30 minutes. Samples were then diluted with five sample volumes of 100 mM ammonium bicarbonate. Trypsin Gold (Promega) was added at a 1:100 (enzyme:protein wt/wt) ratio and lysates were rotated for 16 hours at RT. 10% vol/vol trifluoroacetic acid (TFA) was added to each sample to a final concentration of 0.1% TFA. Samples were desalted under C18 BioPureSPN SPE columns (Nest group) according to the manufacturer’s instructions. Samples were dried by vacuum centrifugation and resuspended in 0.1% formic acid (FA) for mass spectrometry analysis.

### Mass spectrometry data acquisition

All samples were analyzed on an Orbitrap Eclipse mass spectrometry system equipped with an Easy nLC 1200 ultra-high pressure liquid chromatography system interfaced via a Nanospray Flex nanoelectrospray source (Thermo Fisher). Samples were injected onto a fritted fused silica capillary (30 cm × 75 μm inner diameter with a 15 μm tip, CoAnn Technologies) packed with ReprosilPur C18-AQ 1.9 μm particles (Dr. Maisch GmbH). Buffer A consisted of 0.1% Formic Acid (FA), and buffer B consisted of 0.1% FA/80% Acetonitrile (ACN). Peptides were separated by an organic gradient from 5% to 35% mobile buffer B over 120 min followed by an increase to 100% B over 10 min at a flow rate of 300 nL/min. Analytical columns were equilibrated with 3 μL of buffer A. To build a spectral library, samples from each set of biological replicates were pooled and acquired in data-dependent manner. Data-dependent acquisition (DDA) was performed by acquiring a full scan over a m/z range of 375-1025 in the Orbitrap at 120,000 resolving power (@ 200 m/z) with a normalized AGC target of 100%, an RF lens setting of 30%, and an instrument-controlled ion injection time. Dynamic exclusion was set to 30 seconds, with a 10 p.p.m. exclusion width setting. Peptides with charge states 2-6 were selected for MS/MS interrogation using higher energy collisional dissociation (HCD) with a normalized HCD collision energy of 28%, with 3 seconds of MS/MS scans per cycle. Data-independent analysis (DIA) was performed on all individual samples. A full scan was collected at 60,000 resolving power over a scan range of 390-1010 m/z, an instrument controlled AGC target, an RF lens setting of 30%, and an instrument controlled maximum injection time, followed by DIA scans using 8 m/z isolation windows over 400-1000 m/z at a normalized HCD collision energy of 28%.

### Mass spectrometry data analysis

The Spectronaut algorithm was used to build spectral libraries from DDA data, identify peptides/proteins, localize phosphorylation sites, and extract intensity information from DIA data^88^. DDA data were searched against the Homo sapiens reference proteome sequences in the UniProt database (one protein sequence per gene, downloaded on August 23, 2023). False discovery rates were estimated using a decoy database strategy^89^. All data were filtered to achieve a false discovery rate of 0.01 for peptide-spectrum matches, peptide identifications, and protein identifications. Search parameters included a fixed modification for carbamidomethyl cysteine and variable modifications for N-terminal protein acetylation and methionine oxidation. All other search parameters were Biognosys factory defaults. Statistical analysis of proteomics data was conducted utilizing the MSstats package in R. All data were normalized by equalizing median intensities, the summary method was Tukey’s median polish, and the maximum quantile for deciding censored missing values was 0.999. For protein abundance analyses, only the top 10 peptide features per protein were considered.

### Gene ontology enrichment analysis

Gene Ontology (GO) enrichment analysis was performed on proteomics data using a hypergeometric test with the dhyper function in R^90^. Gene ontologies annotations were downloaded from UniProt and GO definitions from the Gene Ontology Resource on Feb. 18, 2021^91,92^. The test sets were comprised of proteins significantly increasing or decreasing (i.e., |log_2_fold-change| > 1 and adjusted p-value < 0.05, excluding infinity values) in each comparison of interest, and the background set was all proteins quantified in the comparison of interest. Enrichment tests were performed for any GO term that had at least 2 overlapping proteins in the test set. Proteins identified by peptides that were not unique to a single protein sequence were excluded from this analysis.

### HIV-PDX memory CD4^+^ T-cell mouse model

All animal procedures and experiments were performed according to NIH regulations and standards on the humane care and use of laboratory animals. Protocols were approved by the Weill Cornell Medical College Institutional Animal Care and Use Committee (protocol 2018-0027). 6-wk-old NSG mice (The Jackson Laboratory) were engrafted with memory CD4^+^ T-cells (5–7 x 10^6^ cells in 100 μl Hank’s Balanced Salt Solution (HBSS)) via tail vein injection. Once memory CD4+ T-cells exceeded 50 cells/μl, mice were engrafted with autologous memory CD8⁺ T-cells (5-7 x 10^6^ cells in 100 μl HBSS) via tail vein injection and simultaneous infection with 10,000 TCID50 HIV_JRCSF_ (in 100 μl media) via i.p. injection. For experiments assessing ARV suppression, mice were administered a cocktail of ARVs (57 mg/kg TDF, 143 mg/kg FTC, and 7 mg/kg DTG) via daily s.c. injections for the indicated duration. Mice were maintained by trained research animal technicians and received daily wellness checks.

For weekly peripheral blood collection, up to 120 μl of blood was collected into EDTA-coated tubes via a tail vein nick technique and processed immediately. Blood was centrifuged at 5,000 RCF for 5 min to separate plasma, which was stored at −80°C before viral RNA extraction. Whole blood was immediately stained for flow cytometric analysis with combinations of the following human specific antibodies: CD4 (BV421, RPA-T4,Biolegend), CD4 (APC-Cy7, A161A1, Biolegend), CD8 (BV605, SK1, Biolegend), CD8 (BUV395, RPA-T8, BD), CD127 (BV510,A019D5,Biolegend), OX40 (BV711, ACT35,Biolegend), CD3 (BV786, OKT3, Biolegend), PD-1 (AF488, EH12.2H7, Biolegend), PD-1(AF700, EH12.2H7, Biolegend), CD45 (PerCP/Cy5.5, QA17A19, Biolegned), anti-HLA-A/B/C (APC-Cy7, W6/32, Biolegend), BIM (AF488, C43C5, Cell Signaling Technology); TOX (PE, REA473, Miltenyi Biotec); H3K27 (PE-Cy7, C3B11, Cell Signaling Technology); TCF-1 (AF647, 7F11A10, Biolegend), TCF-1 (AF647, S33-966, BD), CD95 (BUV737, DX2, BD), CD45RA (BV605, HI100, Biolegend), anti-CD27 (Pe-Dazzle594, M-T271, Biolegend), anti-CCR7 (PE-Cy5, G043H7, Biolegend), Brilliant Violet 711 anti-TNFα,(BV711, MAb11, Biolegend), BCL-2 (AF488, 100, Biolegend), anti-ID2 (Pe-Cy7, ILCID2 eBioscience). SL9-PE and IV9-APC (or IV9-BV421) tetramers (see Tetramer assembly) were included, however as frequencies of specific cells were found to be low no resulting analyses are presented. Live/Dead Aqua (Invitrogen) or Live/Dead Blue, were used as viability markers. Cells were fixed with 4%PFA, or fixed and permeabilized with the Foxp3 / Transcription Factor Staining Buffer Set (eBioscience). Samples were analyzed on an the Cytek Aurora (Cytek Biosciences) or Attune NxT flow cytometer flow cytometer. Data were analyzed using Flowjo software (TreeStar). Cell counts were calculated with CountBrite Absolute counting beads (Invitrogen).

### Fluorescently labeled tetramer assembly

0.4 ug of biotinylated SL9 (SLYNTAVTL) or IV9 (ILKEPVHGV) peptides (NIH Tetramer Core Facility) were conjugated to PE-streptavidin (Agilent), APC-streptavidin (Agilent) or BV421-streptavidin (BD). Tetramers were assembled by the stepwise addition of fluorophore-conjugated streptavidin to a final biotin-peptide: streptavidin ratio of 4:1 over 10 incubation intervals of 7-10 minutes at room temperature.

### Analysis of HIV viral load

HIV infection in mice was monitored weekly by measuring HIV RNA concentrations in plasma using the previously described integrase single-copy assay protocol (Cillo et al., 2014). Plasma RNA was isolated with the QIAamp viral RNA mini kit. Samples were analyzed on QuantStudio 6 flex and QuantStudio 7 Pro Real-Time PCR Systems using the following cycling parameters: 45°C for 10 min, 95°C for 10 min, followed by 40 cycles of 95°C for 15 s and 60°C for 1 min. Cycle threshold values were compared with a validated HIV RNA standard run on each plate to determine HIV RNA concentration.

### Isolation of mouse spleen and bone marrow cells

Mice were humanely euthanized by inhalation of CO_2_ anesthesia. The spleen was collected and homogenized in R10 containing 25 U/mL DNAase. Bone marrow cells were isolated by making a superficial cut on the tip of the isolated femur and tibia followed by centrifugation at 15,000 x g for 10 minutes. Red blood cells were lysed with ACK buffer at 37°C for 5 min. Cells were viably frozen. or prepared as dry pellet and stored in -80°C for DNA and cell-associated RNA extraction.

### Ex vivo stimulation of spleen cells for TCF-1 expression analysis and IFN-γ quantification

5 x 10^6^ cells from each mouse spleen were thawed and rested for 3 hours at 37 °C 5% CO2 in R10 medium. Cells were then divided into seven conditions in triplicate with 700,000–1,000,000 cells per well per condition and stimulated for 16 hours with the following whole gene product peptide pools from the NIH HIV Reagent Program at 1ug/mL: HIV-Gag (ARP-12425), HIV-Pol (ARP-12438), HIV-Env (ARP-190434) and HIV-Nef (ARP-12545); from JPT Peptide Technologies. Phytohemagglutinin (PHA) was added at 2ug/mL as a positive control, and 0.5% DMSO in R10 medium was used as a negative control. 1 μl of anti-CD107a (LAMP-1) Antibody (BV785, H4A3, Biolegened) was added to each well. Post-stimulation, cell supernatant was collected and frozen at -80C. Cells were washed in 2% FBS 2 mM EDTA-PBS and surface stained with the following anti-human antibodies: CD45 (PerCP/Cyanine5.5, 2D1, Biolegend), CD137 (BV650, 4-1BB, Biolegend), CD8 (BV605, SK1, Biolegend), TOX (PE, REA473, Miltenyi), CD134 (OX40) (BV711, Ber-ACT35, Biolegend), PD-1 (AF488, EH12.2H7, Biolegend), TCF1 (AF647, 7F11A10, Biolegend), CD69 (PE-eFluor 610, FN50 Thermo Fisher), HLA-A,B,C (APC/Cy7, W6/32, Biolegend), CD4 (BV421, RPA-T4, RPA-T4). Live/Dead Aqua Fixable cell stain (Invitrogen) was used as a viability marker. Cells were fixed and permeabilized with the Foxp3/Transcription Factor Staining Buffer Set (eBioscience). Samples were analyzed on an Attune NxT flow cytometer and Flowjo software (TreeStar).

### IFN-γ ELISA

Concentrations of IFN-γ in mouse plasma and ex vivo stimulation supernatants were determined using the ELISA MAX Deluxe Set Human IFN-γ Kit (BioLegend, cat #430104) according to the manufacturer’s recommendations. Plasma was diluted 1:1 or 1:10 in 1× Assay Diluent A. Optical density was read at 450 nm and 570 nm using a SpectraMax i3x platform (Molecular Devices).

### Cell-associated HIV RNA

DNA and RNA were co-extracted from isolated CD4+ cells using the AllPrep Mini kit (Qiagen, cat #80204). RNA was used for total polyadenylated cDNA generation using dT20 primer and reverse transcription was performed with SuperScript IV First-Strand Synthesis (Thermo Fisher, 18091150). Resulting HIV-cDNA levels were quantified using BIO-RAD QX200 droplet digital PCR using the following HIV-specific primer and probe sets targeting two regions of the viral transcriptome: HXB2 coordinates 684-810 for unspliced HIV mRNA (Forward primer: 5’- TCTCGACGCAGGACTCG-3’, reverse primer 5’-TACTGACGCTCTCGCACC-3’, and probe 5’- /56-FAM/CTCTCTCCT/ ZEN/TCTAGCCTC/31ABkFQ/−3’); HXB2 coordinates 9435-9525 for total polyadenylated viral RNA with forward primer 5’- GGGACTTTCCGCTGGG-3’, reverse primer 5’- AGCAGCTGCTTA- TATGCAG-3’, and probe 5’-/56-FAM/TGAGGGCTC/ZEN/GCCACTCC/ 3IABkFQ/−3’. DdPCR data analyses were performed using the BIO-RAD QuantaSoft software suite.

### Cell-associated HIV DNA (IPDA)

Genomic DNA was isolated from splenocytes using the QIAamp DNA Mini Kit (Qiagen) with precautions to minimize DNA shearing. Intact HIV-1 copies/million CD4^+^ T-cells were determined by droplet digital PCR (ddPCR) using the Intact Proviral DNA Assay (IPDA), where HIV-1 and human RPP30 reactions were conducted independently in parallel, and copies were normalized to the quantity of input DNA and %CD4^+^ cells in starting material (as determined by flow cytometry). In each ddPCR reaction, genomic DNA was combined with ddPCR Supermix for Probes (no dUTPs, BioRad), primers, probes and nuclease free water. Primer and probe sequences (5′–>3′) were: RPP30 Forward Primer- GATTTGGACCTGCGAGCG; RPP30 Probe- VIC-CTGACCTGAAGGCTCT-MGBNFQ; RPP30 Reverse Primer-GCGGCTGTCTCCACAAGT; RPP30-Shear Forward Primer- CCATTTGCTGCTCCTTGGG; RPP30-Shear Probe- FAM- AAGGAGCAAGGTTCTATTGTAG-MGBNFQ; RPP30-Shear Reverse Primer- CATGCAAAGGAGGAAGCCG;HIV-1 Ψ Forward Primer- CAGGACTCGGCTTGCTGAAG; HIV-1 Ψ Probe- FAM- TTTTGGCGTACTCACCAGT- MGBNFQ; HIV-1 Ψ Reverse Primer- GCACCCATCTCTCTCCTTCTAGC; HIV-1 env Forward Primer- AGTGGTGCAGAGAGAAAAAAGAGC or HIV-1 env Modified Forward Primer – GTCTGGCCTGTACCGTCAGT; HIV-1 env Probe- VIC-CCTTGGGTTCTTGGGA-MGBNFQ; HIV- 1 env Reverse Primer- GTCTGGCCTGTACCGTCAGC. Droplets were prepared using either the Automated Droplet Generator (BioRad) and cycled at 95 °C for 10 min; 45 cycles of (94 °C for 30 sec, 59 °C for 1 min) and 98 °C for 10 min. Droplets were analyzed on a QX600 Droplet Reader (BioRad) using QuantaSoft software (BioRad, version 1.7.4). Two technical replicates were performed for each sample. Intact HIV-1 copies (Ψ and env double-positive droplets) were corrected for DNA shearing based on the frequency of RPP30 and RPP30-Shear double-positive droplets. Limit of Detection (LOD) was determined for each sample based on the input human genome (RPP30) equivalents assayed per reaction with correction for DNA shearing index (DSI). We confirmed that these RPP30 primers and probes do not amplify a target in mouse DNA.

### Ultrasensitive p24 ELISA

P24 plasma concentrations were quantified as previously described^93^. Briefly, anchor antibody microprinted microwell plates were incubated with an anti-p24 peptide-tagged capture. Inactivated (1% Triton) mouse plasma was diluted 1:4 in assay diluent and incubated directly on plates. Bound p24 antigen is then sandwiched with a p24 biotinylated detector antibody. Streptavidin horseradish peroxidase, chemiluminescent substrate Luminol and peroxide is used for detection. Chemiluminescence signal is scanned on the SP-X imager (Quanterix). The data file was analyzed using the Quanterix SP-X Analysis Software V2.1.1.7737 where a standard curve using 5 parametric logarithmic (5PL) curve fit %CV, LOD, LLOQ, and R^2^ were created. Plasma from naïve mice were used to confirm the assay limit of detection (LOD).

### IFN-γ ELISPOT

HIV IFN-γ ELISPOT assays were performed following the manufacturer’s instructions (Mabtech ELISPOT Flex Human IFN-γ). In brief, MultiScreen IP 96-well plates (Millipore) were coated with 15 µg/mL of anti-IFN-γ antibody (clone 1-D1K) in phosphate-buffered saline and incubated overnight at 4°C. Plates were washed 5x, peripheral blood mononuclear cells were added at 2 × 10^5^ cells per well with HIV (Gag, Pol, Nef, Tat, Rev, Vpr, Vpu, Vif) or CMV-pp65 peptide pools (1 µg/mL/peptide) and Staphylococcal enterotoxin B (1 µg/mL, Toxin Technologies) or DMSO control (0.5% v/v). Plates were incubated overnight, washed 5x and secondary antibody was added at 1 µg/mL (clone 7-B6-1) and incubated for 2 hours. Plates were washed 5x again and incubated with Streptavidin-ALP (Mabtech) for 1 hour, then washed 5x and developed with Color Development buffer (Bio-Rad) for 15 minutes. Plates were washed, dried overnight and spots were counted with the ImmunoSpot instrument (C.T.L). ELISPOT responses against whole gene product peptide pools were background subtracted, but no other positivity criterion was applied.

## Data availability

Data that support the findings of the present study are available upon request via email to the lead corresponding author R.B.J. Data involving human research participants are subject to the data protection constraints in the written informed consent signed by the study participants.

## Acknowledgements

We thank all study participants who devoted time to our research as well as the clinical research team involved in the study. We thank the NIH Tetramer Core Facility (contract number 75N93020D00005) for providing biotinylated SL9 and IV9 peptide monomers used to identify HIV-specific cells. We thank Afam Okoye and Benjamin Varco-Merth for providing the ART supply used for in vivo experiments. We thank Ari Melnick, Wendy Béguelin, and Yusuke Isshiki for guidance on tazemetostat resuspension. We thank Guinevere Q. Lee and Nelson Sonela for ddPCR support. This work was supported by the following grants: R01MH130197, R01AI170239, R37AI181626, R01AI176601, R01AI176943, R01AI170245, R01AI165031, UM1AI164562, UM1AI164565, R01AI150412, R21AI170246, R01AI147845, U01AI145921, R01AI167691, R21AI172554. The funders had no role in study design, data collection and analysis, decision to publish or preparation of the manuscript. Figures and figure schematics were created using GraphPad Prism, BioRender.com and Adobe Illustrator.

## Contributions

A.G. and R.B.J. designed the research; A.G., IGM., A.D, F.K., T.J.K., T.T.H., L.L., Y.R., T.K., D.C.C., U.C., C.L., S.T., T.M., P.Z., performed experimental work in vitro; A.D coordinated and supervised the animal experiments; I.G.M., J.W., D.C.J., T.L.D., E.F.I performed the experimental work in vivo; S.B and K.L.C contributed the custom JRCSF (ΔNef) infectious molecular clone. A.G., I.G.M, A.R.W., A.D., M.P, R.B.J participated in discussion and interpretation of the results. A.G and A.R.W performed the statistical analyses. A.G., I.G.M. and R.B.J. prepared figures and wrote the manuscript. All authors reviewed the manuscript and approved it for publication. R.B.J., K.L.C., A.B., L.M.E., C.L, D.B, J.R.J supervised experiments and analyzes.

## Extended Data

**Extended Data Fig. 1:**
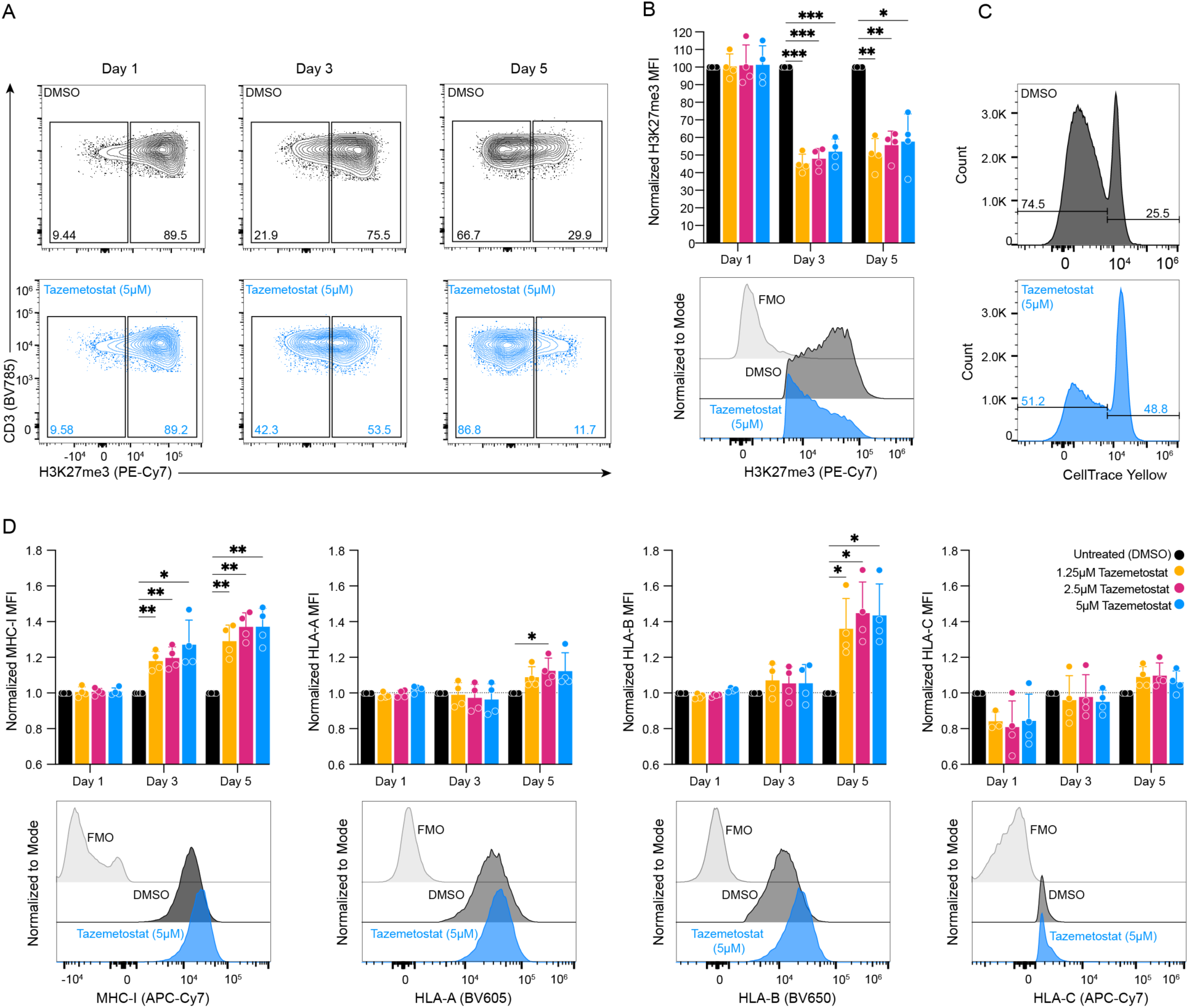
Treatment with tazemetostat reduces H3K27 tri-methylation, decreases proliferation, and increases HLA-B surface levels on CD4^+^ T-cells. Uninfected CD4^+^ T-cells were activated in the presence of anti- CD3, anti-CD28 antibodies and then treated for 1, 3 or 5 days with increasing concentrations of tazemetostat. Results are color-coded as follows: black = DMSO (untreated control), yellow = 1.25μM, red = 2.5μM, blue = 5μM. **a**) Representative flow cytometry plots, gated on viable CD4^+^ T-cells and showing % of H3K27me3^+^ and H3K27me3^-^ cells that were either left untreated (black, top row) or treated with 5 μM tazememtostat (blue, bottom row) at the indicated time-point. **b**) Top panel: Summary flow cytometry data from 4 different donors. Shown are means ± SD, normalized to the geomean MFI of the untreated condition at each timepoint. Lower panel, overlay of results from one donor a with fluorescence-minus-one (FMO) control (gray). **c**) Cells were labeled with CellTrace Yellow to enable assessment of proliferation (by dye diminution). Shown are histograms following 5 days treatment with DMSO (black, top) or tazemetostat (blue, bottom), showing that tazemetostat impaired proliferation. These correspond to the samples in panel a. **d**) Cell surface MHC-I (HLA-A,B,C), HLA-A, HLA-B and HLA-C in cells treated with the indicated concentrations of tazemetostat, for 1, 3 or 5 days. Top panels: Shown are flow cytometry data from the same experiment in panel b (4 donors without HIV) means ± SD. Bottom panels: Respective fluorescence minus one (FMO) controls (gray) and fluorescence intensities of representative samples either left untreated (black) or treated with 5 μM tazememtostat (blue).

**Extended Data Fig. 2:**
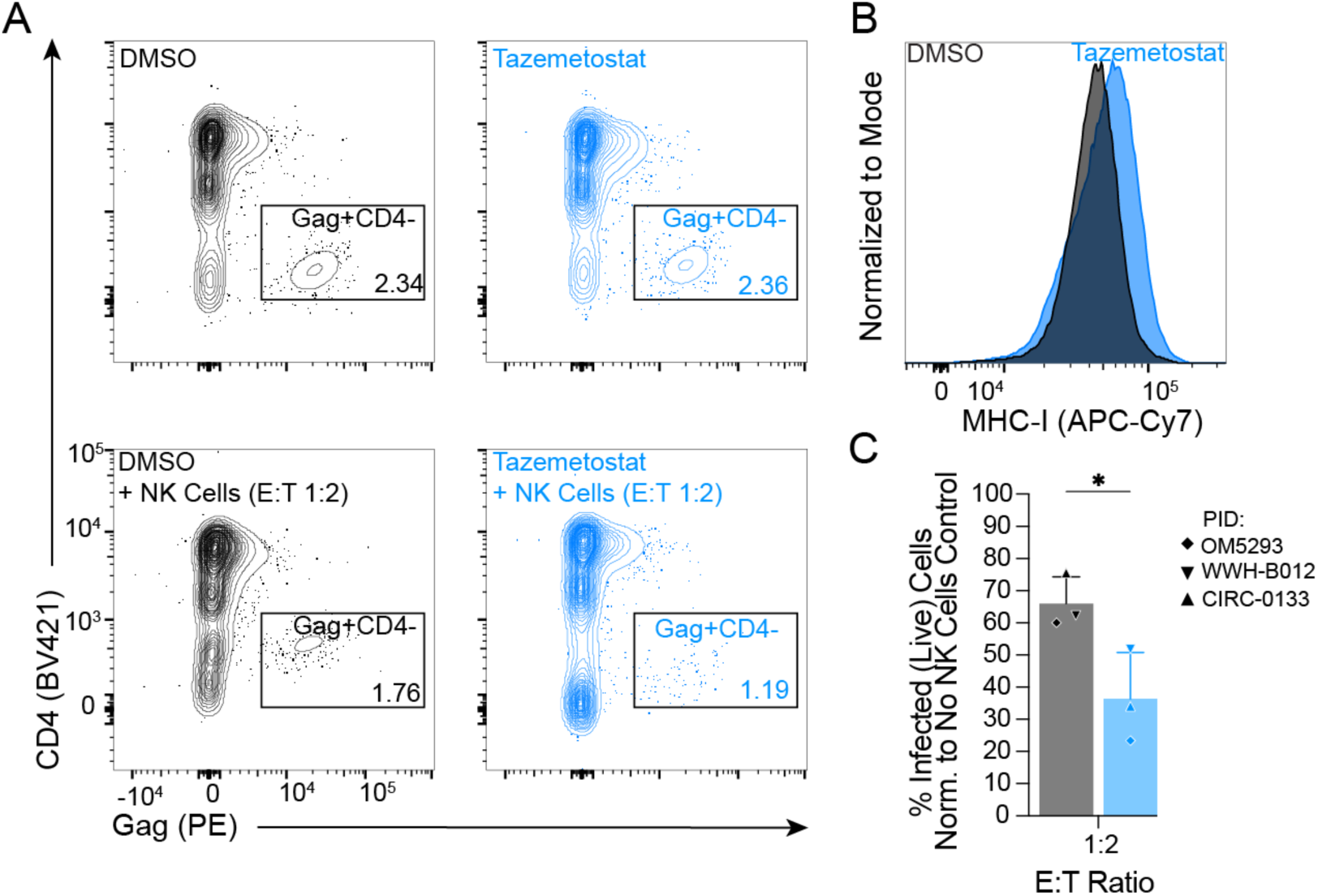
Tazemetostat and enhances NK-mediated elimination of infected cells in vitro. **a**) Flow cytometry plots depicting viable CD4^+^ T-cells treated with 5 μM tazemetostat (blue), or DMSO (black), alone or after co-culture with an autologous NK-cells. **b**) Representative histogram depicting MHC-I fluorescence intensity of CD4^+^ T-cells treated with 5μM tazemetostat (blue), or DMSO (black). **c**) % infected CD4^+^ T-cells (within the Live-cell gate) isolated from 3 PWH (refer to Method section for clinical information on study participants) after treatment with tazemetostat (blue), or DMSO (black) and exposure to autologous NK-cells at 1:2 effector:target (E:T) ratio. Values are shape-coded based on participant ID (PID). Shown are Mean and SD. Statistical significance was determined by 2- way ANOVA. **P < 0.01, ***P < 0.001. ****P<0.0001.

**Extended Data Fig. 3:**
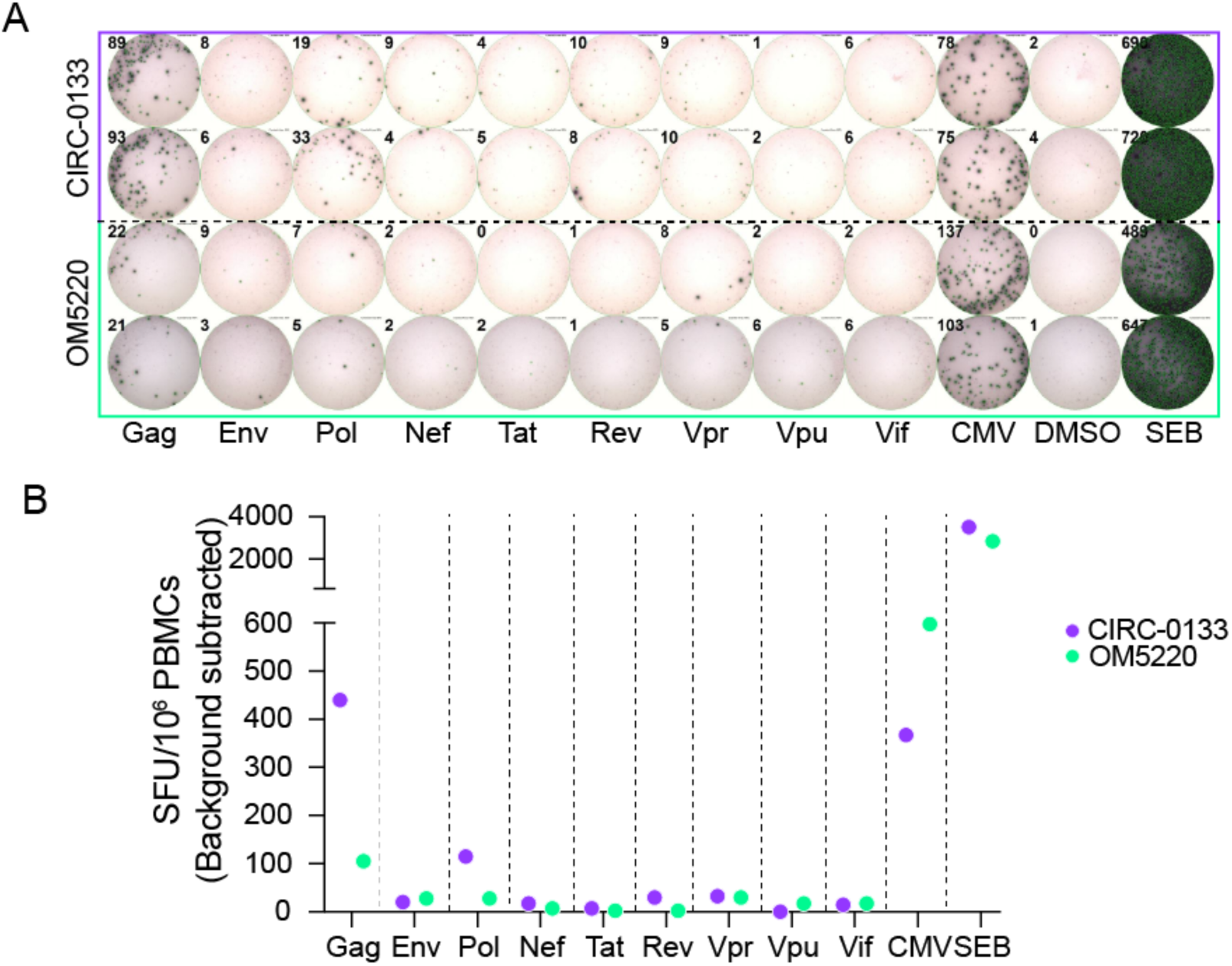
Magnitudes of IFN-γ producing HIV-specific T-cell responses. **a)** Representative IFN-γ enzyme-linked immune absorbent spot (ELISPOT) results for 2 participants (CIRC-0133, top and OM5220, bottom) with 2 × 10^5 PBMCs/well, in two technical replicates. Dotted line represents separation between samples of the two donors. **b**) Magnitudes of IFN-γ responses calculated from a. Each data point represents the mean SFU/10^6 PBMCs following background subtraction of negative control wells (duplicates).

**Extended Data Fig. 4:**
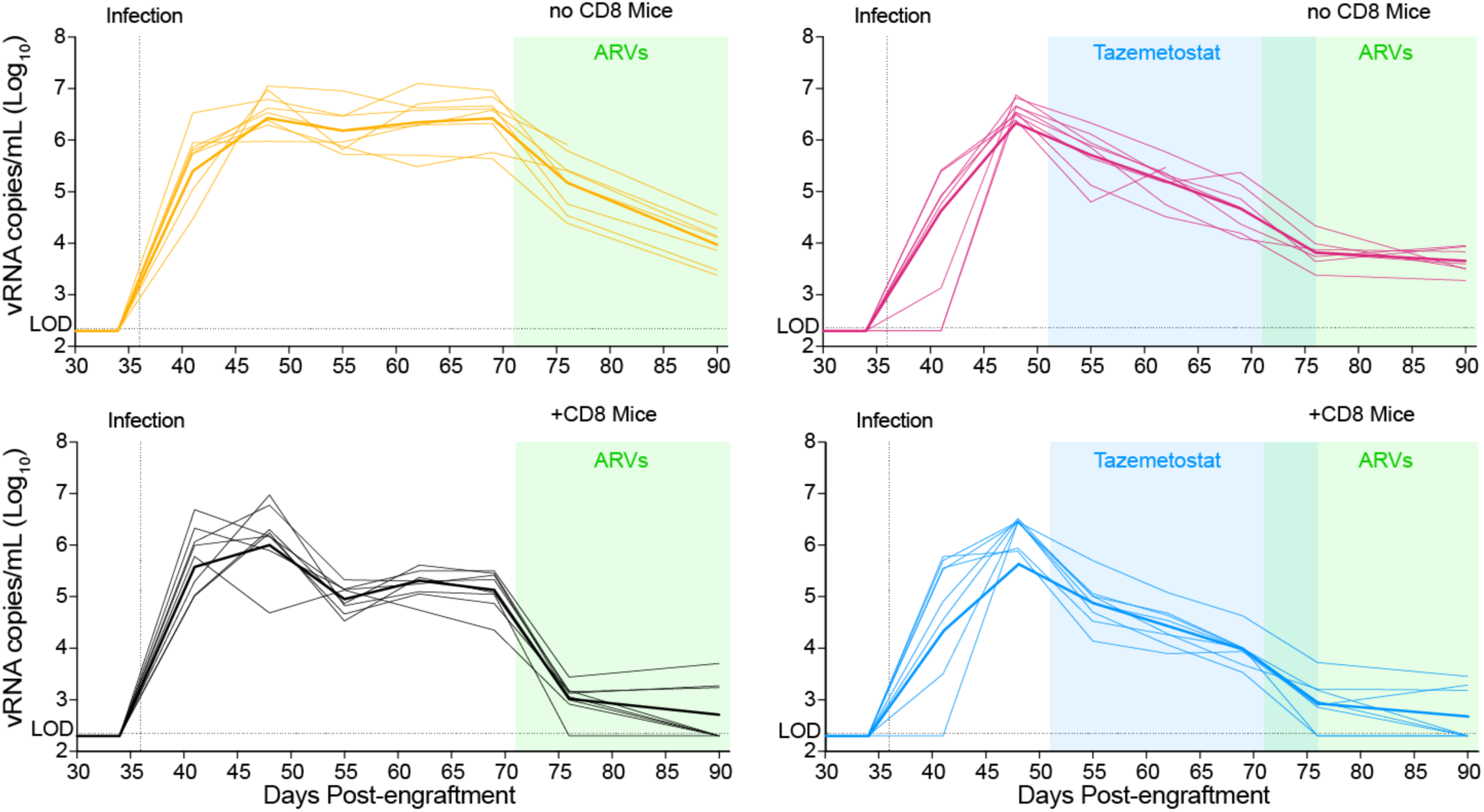
Longitudinal quantification of HIV plasma viral load in vehicle-treated mice engrafted with CD4^+^ T-cells only (“no CD8” mice, yellow), tazemetostat-treated mice engrafted with CD4^+^ T-cells only (“no CD8” mice, magenta), vehicle-treated mice engrafted with CD4^+^ and CD8^+^ T-cells (“+CD8” mice, black), and tazemetostat-treated mice engrafted with CD4^+^ and CD8^+^ T-cells (“+CD8” mice, blue). Thin lines represent viral load of single mice, thick lines represent median values of mice of the same group. Blue and green rectangles in b and c mark the timeframe of tazemetostat administration (day 50 - 76) and ARVs administration (day 71 - 107), respectively.

**Extended Data Fig. 5:**
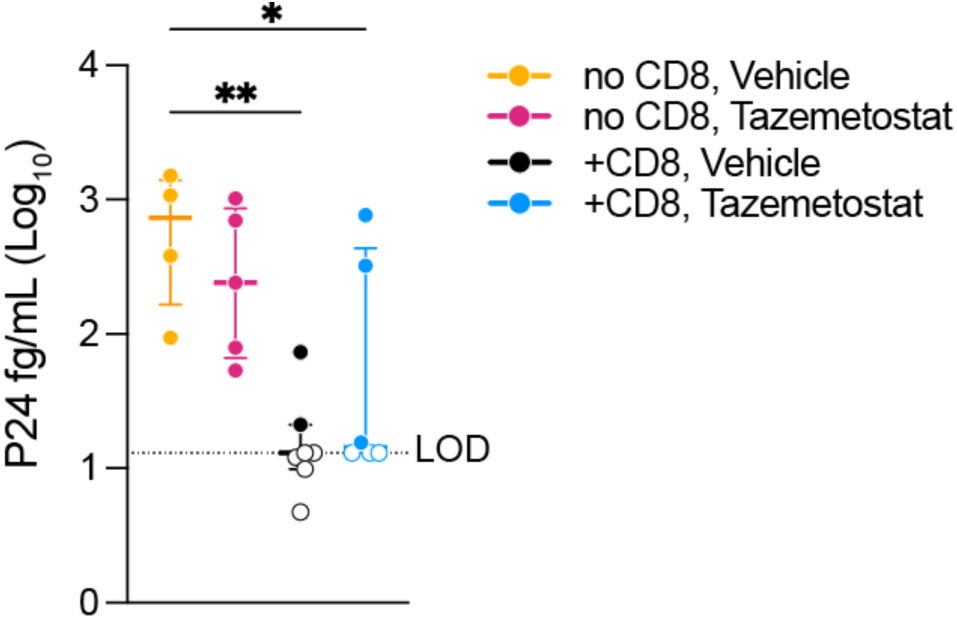
HIV-p24 antigen quantification in mouse plasma at day 163. No CD8, Vehicle group in yellow, No CD8, tazemetostat group in magenta, +CD8, Vehicle group in black, +CD8, tazemetostat group in blue. Each dot represents a single mouse. Open circles indicate value at or below the limit of detection (13.04 fg/mL). Median values with interquartile range are shown. Statistical significance was determined by one-way ANOVA with multiple comparison adjustment. *P <0.05, **P<0.01.

**Extended Data Fig. 6:**
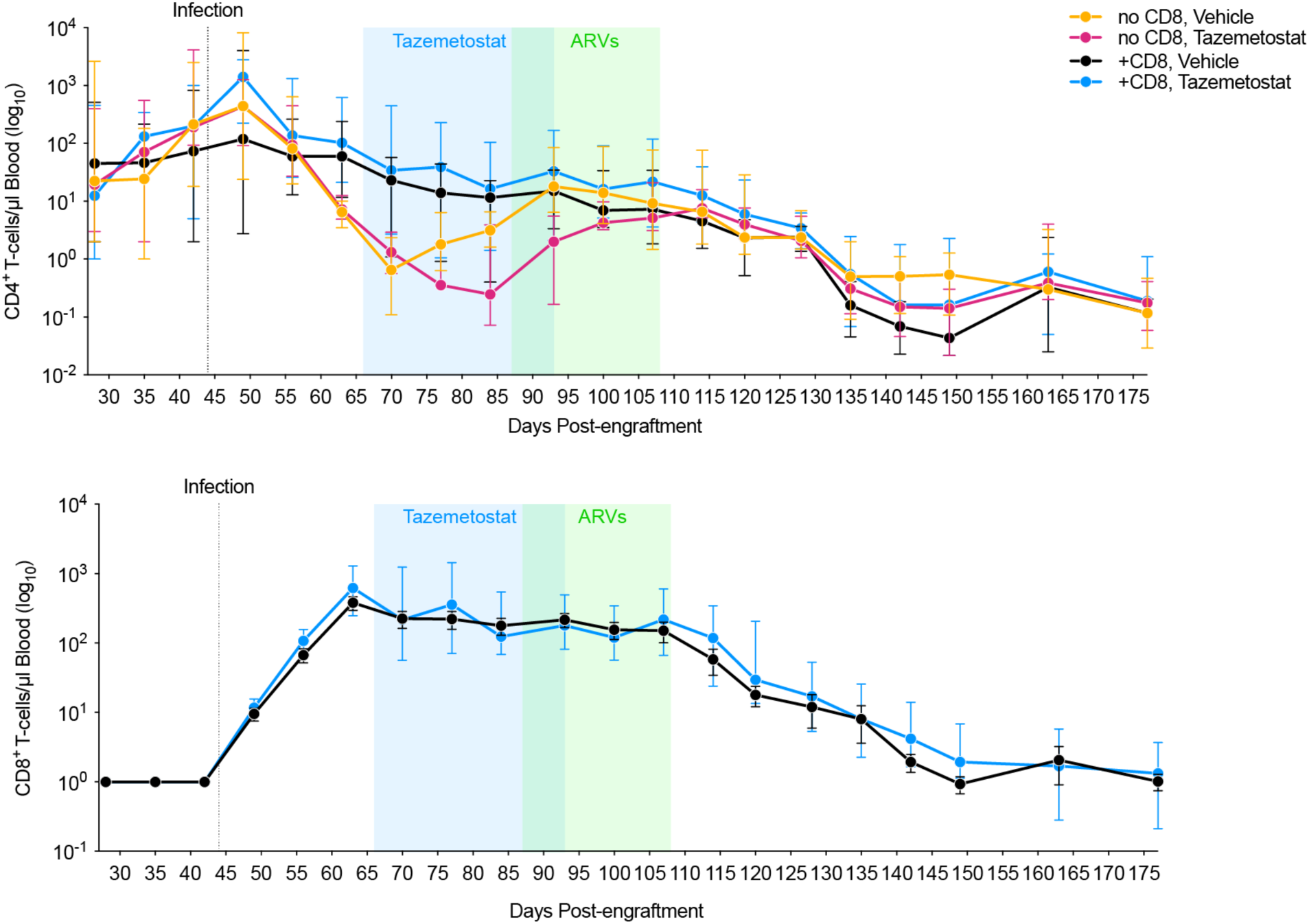
Longitudinal quantification of CD4^+^ and CD8^+^ T-cell counts in vivo. At each time-point (i.e., weekly bleed) median values with interquartile range are shown.

**Extended Data Fig. 7:**
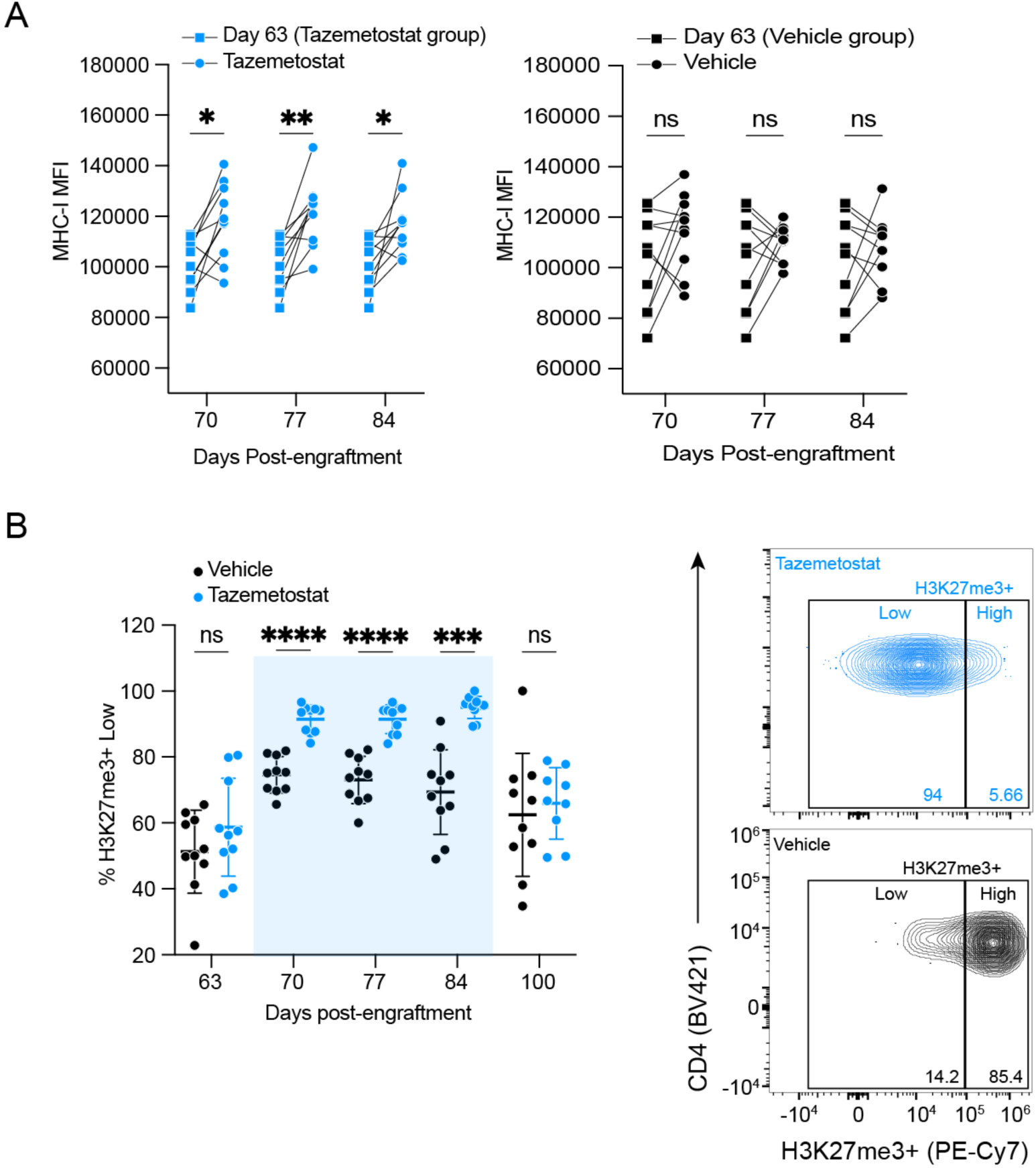
**a**) Left: Changes in MHC-I MFI (geomean) in tazemetostat-treated mice before (day 63) and during (days 70, 77, 84) treatment. Right: Changes in MHC-I MFI (geomean) in Vehicle-treated mice before (day 63) and during (days 70, 77, 84) treatment. **b**) Left: Longitudinal changes in % of H3K27me3^+^ “Low” CD4^+^ T-cells in tazemetostat-treated (blue) or Vehicle-treated (black) mice at the indicated time-points. Each dot represents a single mouse. Shown are Mean ± SD. Statistical significance was determined by 2-way ANOVA with multiple comparison adjustment. ***P < 0.001. Blue rectangle marks time period of tazemetostat administration (day 66 - 87). Right: Representative flow cytometry plots, gated on CD4+ T-cells and showing % of H3K27me3^+^ “High” and “Low” cells from either a tazemetostat-treated (blue) or Vehicle-treated (black) mouse at day 77 post-engraftment.

**Extended Data Fig. 8:**
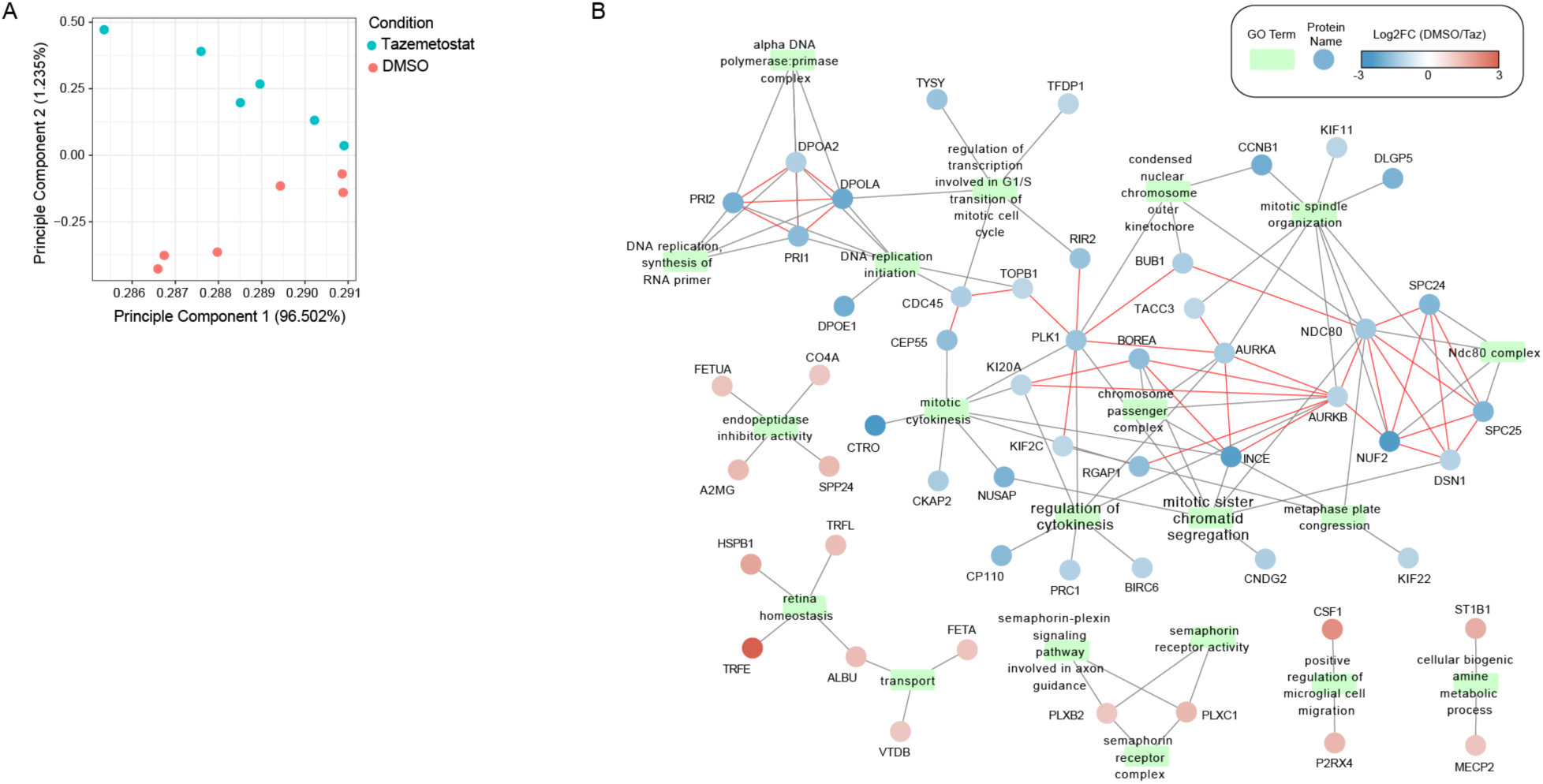
**a**) Principal component analysis (PCA) of protein expression profiles for DMSO and tazemetostat-treated samples. Each point represents an individual sample, colored by condition (blue: tazemetostat, red: DMSO). The first two principal components account for 58.302% and 12.35% of the total variance, respectively, with clear separation between the two conditions. **b**) Network view of the GO terms and the proteins that are driving those enrichments. In the network figure, the black edges are proteins associated with GO terms and red edges are protein-protein interactions in the BioGRID multivalidated data set. The color of the nodes reflects the log2fold-change of that protein comparing DMSO to drug.

## Supplementary Information

**Supplementary Table 1:** Characteristics of the study participants. . List of abbreviations: Cobicistat (COBI), Darunavir (DRV), Elvitegravir (EVG), Emtricitabine (FTC), Tenofovir alafenamide (TAF), Tenofovir disoproxil (TDF), Lamivudine (3TC), Abacavir (ABC), Rilpivirine (RPV).

| Participant ID | Ethnicity | Sex assigned at birth | Year of HIV diagnosis | Year of ART initiation | Most recent CD4 count (cells/mm<sup>3</sup>) | Nadir CD4 if known (cells/mm<sup>3</sup>) | Current ART regimen |
| --- | --- | --- | --- | --- | --- | --- | --- |
| OM5220 | Asian | Male | 2010 | 2010 | 420 | 370 | 3TC, ABC, DTG |
| OM5267 | Caucasian | Male | 2011 | 2011 | 480 | NA | 3TC, ABC, DTG |
| OM5293 | Caucasian | Male | 2012 | 2012 | 490 | 430 | CPBI, EVG, FTC, TAF |
| CIRC-0133 | Hispanic | Male | 2003 | 2008 | 526 | 507 | 3TC, ABC, DTG |
| WWH-012 | African American | Female | 1997 | 1997 | 968 | 350 | RPV, TAF, FTC |
| WWH-021 | African American | Male | 1998 | 1998 | 874 | 350 | DRV, FTC, TAF |
| WWH-022 | Caucasian | Male | 2012 | 2012 | 1108 | NA | CPBI, EVG, FTC, TAF |

**Supplementary Fig. 1:**
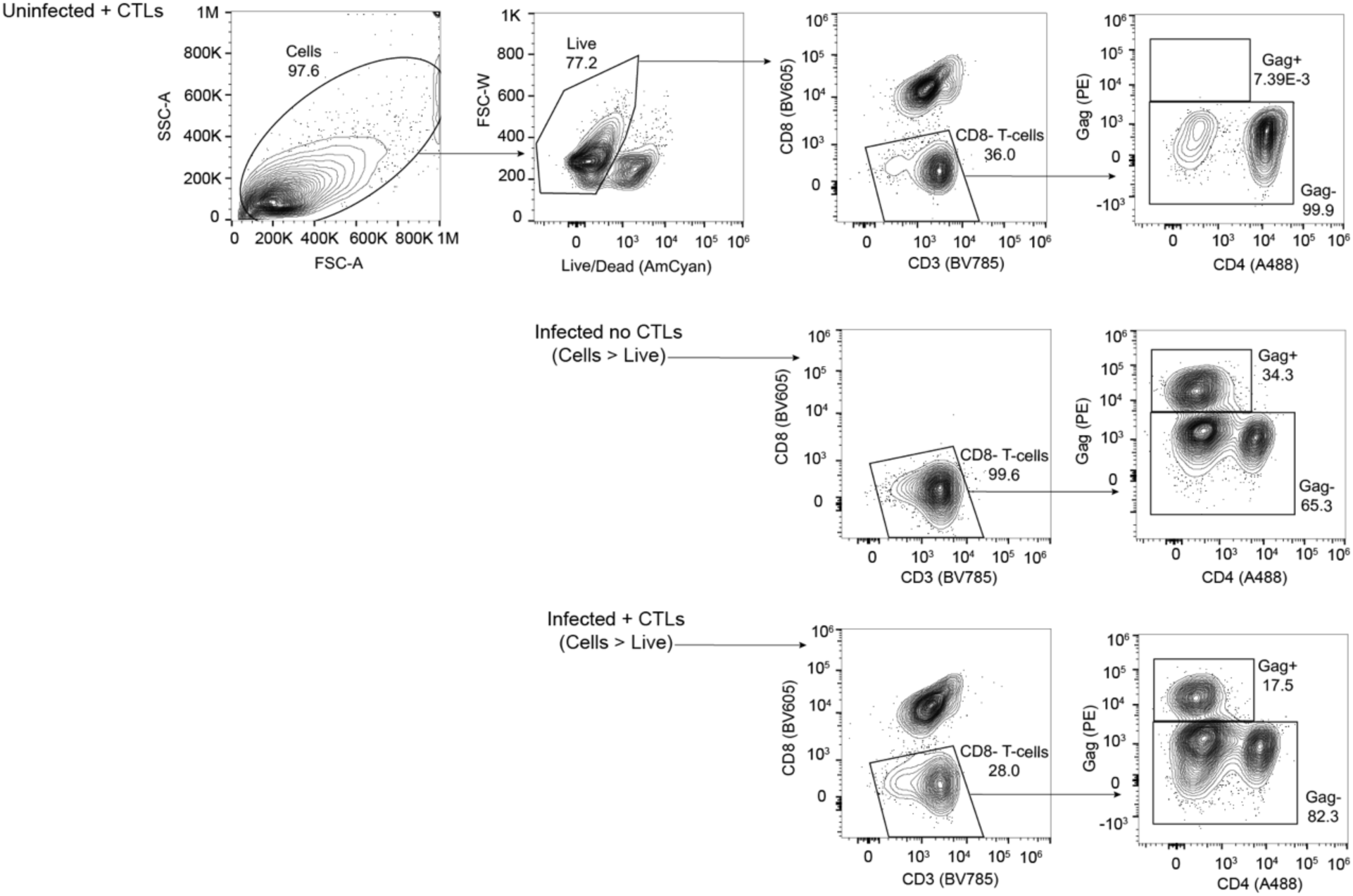
Gating strategy used for flow sorting of infected cells from donor OM5267 and donor OM5220, in the presence (+ CTLs) or absence (no CTLs) of HIV-specific CTL clones.

**Supplementary Fig. 2:**
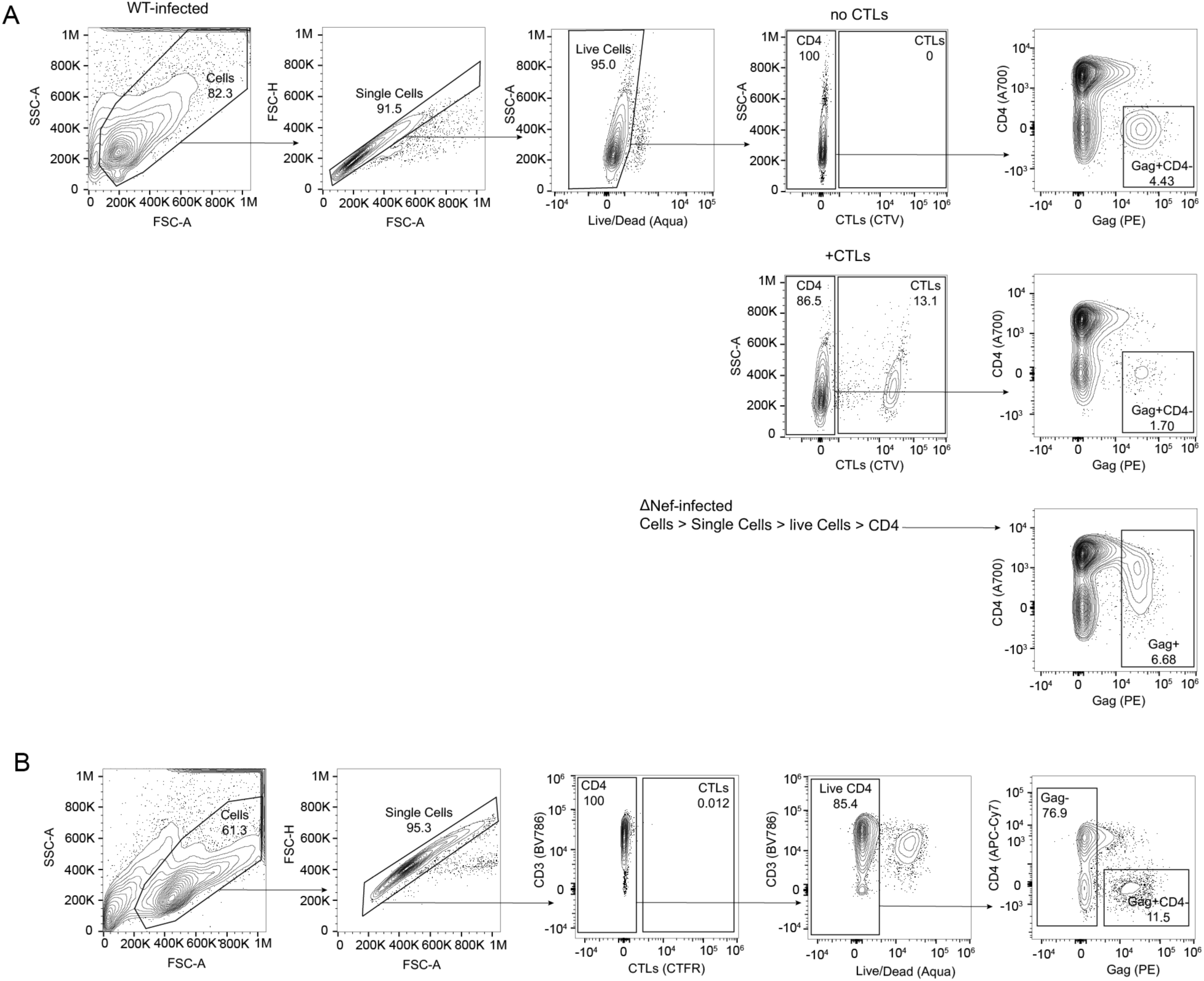
Gating strategy used to assess killing of infected cells in the **a**) in vitro killing assay, (both with WT- and ΔNef-infected cells) and **b**) VIA. ΔNef-infected cells were defined as Gag^+^, CD4^+/-^, taking into consideration the lack of downregulation of CD4 in the absence of HIV-Nef.

**Supplementary Fig. 3:**
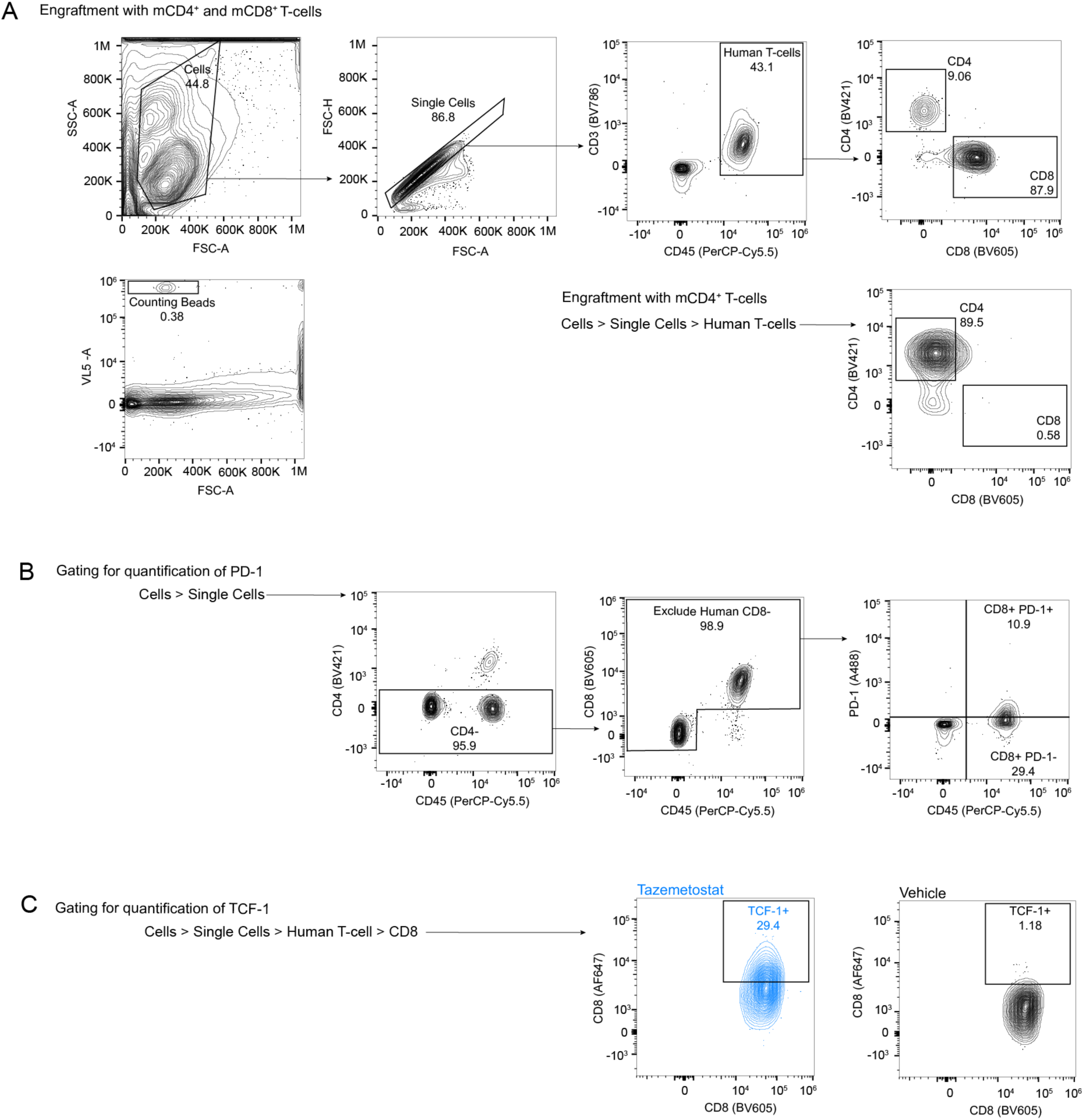
**a**) Gating strategy used for in vivo quantification of CD4^+^ and CD8^+^ T-cell counts. Counting beads were gated based on FSC-A and “VL5”, the Attune NxT detector excited by 488nm laser, filtering emission at 710/50nm. **b - c**) Gating strategy used for in vivo quantification of surface PD-1 and TCF-1, respectively. In c, examples of TCF-1 staining in tazemetostat-treated (blue) and Vehicle-treated (black) mice are shown.

**Supplementary Fig. 4:**
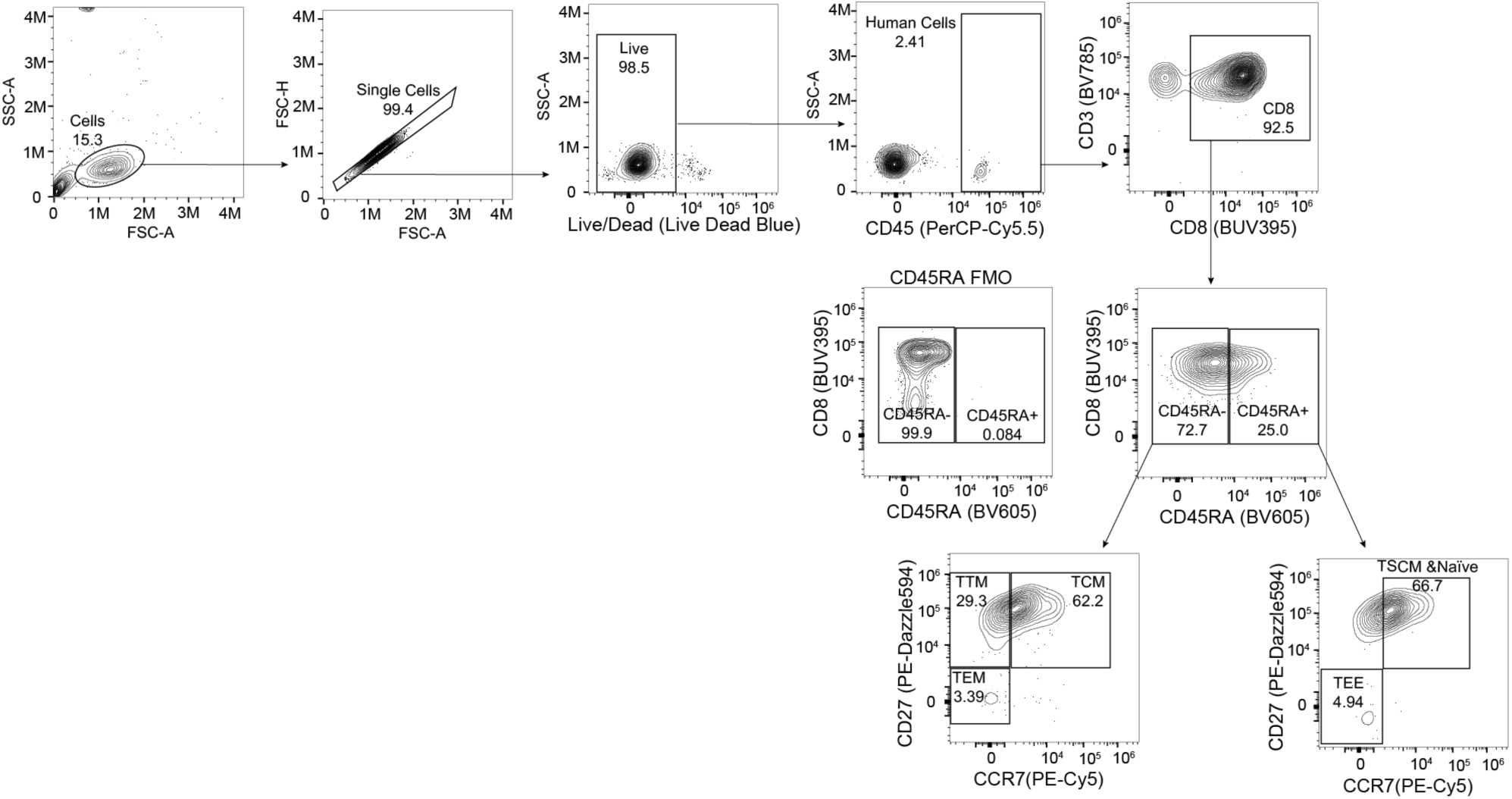
Gating strategy used for in vivo phenotyping of CD8^+^ T-cell sub-populations.

